# Tetraspanin changes in septic blood are organized by cell type, with monocyte CD82 induction and leukocyte CD37 loss associated with mortality

**DOI:** 10.64898/2026.10.14.756292

**Authors:** Jonathan D. Wren, Yingjun Ding, Xin A. Zhang

## Abstract

**Background:** Tetraspanins organize membrane domains that regulate inflammation. The antagonistic pair CD82 (pro-inflammatory, pro-vascular leakage) and CD151 (anti-inflammatory, anti-vascular leakage) coordinate endothelial activation and dysfunction in severe and systemic inflammation such as sepsis. Whether blood-cell tetraspanin networks are reorganized and regulated in cell-type-specific and -coordinated fashions in human sepsis and relate to mortality is unknown.

**Methods:** We integrated bulk meta-analysis (8 ICU-sampled cohorts, 241 controls, 550 sepsis patients), 28-day survival analysis (GSE65682, n=468), single-cell RNA-sequencing (3 datasets, 1.6 million PBMCs from 184 donors), co-expression networks (WGCNA), immune deconvolution with composition-adjusted survival, and mortality prediction models judged against matched random-gene controls. Mendelian randomization tested whether inherited variation in tetraspanin expression alters sepsis susceptibility. Single-cell RNA-sequencing of LPS-challenged mouse lung provided an orthogonal test.

**Results:** Thirteen of 33 tetraspanins changed significantly in septic blood (up: CD63, CD82, CD151, ROM1, TSPAN9, UPK1A; down: CD81, TSPAN3, TSPAN13, TSPAN32, TSPAN18, TSPAN4, TSPAN31), and the direction of change followed myeloid versus lymphoid lineage bias, reflecting the leukocyte composition shift of sepsis. WGCNA placed CD82 and CD151 in distinct modules. The CD82:CD151 ratio did not predict 28-day mortality (HR=1.09, p=0.37). In the discovery cohort, CD37 was the strongest survival-associated tetraspanin (HR=0.72 per SD, FDR=0.002), even after adjusting for leukocyte composition (HR=0.70, p=0.007). Single-cell analysis of three datasets of sepsis patients resolved two cell-type-specific programs, monocyte CD82 up-regulation and pan-leukocyte CD37 loss. CD37 expression was directionally protective in all three independent ICU cohorts, significantly so only in the largest. Mendelian randomization found no evidence that genetically proxied baseline tetraspanin expression affects sepsis susceptibility (0 of 24 genes). In mouse lung, we found macrophage CD82 increased and endothelial CD151 decreased after LPS.

**Conclusions:** Blood-cell tetraspanin networks are reorganized cell-type-specifically in sepsis. Much of the bulk signal reflects leukocyte composition rather than a coordinated membrane program. The CD82/CD151 antagonism, established in endothelium, is not captured by whole-blood ratios, in which the two genes reflect different cell types.

## Introduction

Sepsis is a life-threatening organ dysfunction caused by a dysregulated host response to infection^1^, and a leading global cause of death, with an estimated 48.9 million cases and 11 million deaths annually^2^. Its pathophysiology is defined by endothelial barrier failure and subsequent vascular leakage^3^, which reflects a failure of the normal balance between barrier-disrupting and barrier-stabilizing signals in endothelia, and by coagulopathy, which reflects a dysregulated balance between procoagulant and anticoagulant activity. Tetraspanins are a family of four-transmembrane proteins that laterally associate with partner proteins and with cholesterol and saturated lipids to assemble tetraspanin-enriched membrane domains (TEMDs), which regulate the clustering, trafficking, and endolysosomal turnover of their associated molecules^4^. TEMDs regulate this balance in the endothelium of the vascular wall, yet how they change in circulating blood cells, and whether they are perturbed in sepsis, is unclear.

Our prior work has established that the tetraspanin CD82 reorganizes TEMDs and lipid rafts through cholesterol redistribution and cholesterol-dependent endocytosis, directing partner proteins to late endosomes and lysosomes^5^. In the vasculature, CD82 restrains pathological angiogenesis by limiting GM1-containing lipid raft clustering and the surface presentation of cell-adhesion molecules such as CD44 in endothelial cells^6^. More recently, endothelial CD82 was shown to promote vascular leakage during inflammation by coalescing and retaining accessible cholesterol at the plasma membrane via oxysterol-binding-protein-related protein (ORP)-mediated non-vesicular transport, to tip the balance of antagonistic Rho-family GTPases (confining Cdc42 while maintaining RhoA activity), and statin or ORP inhibition reverses this effect^7^. CD82 therefore behaves as a pro-inflammatory, vascular leakage-facilitating tetraspanin.

The closely related tetraspanin CD151 acts in the opposite direction. CD151 maintains vascular stability and limits permeability by reinforcing endothelial cell-cell and cell-matrix adhesion and balancing RhoA and Rac1 activities^8^. At the subcellular level, CD151 sustains endolysosomal protein quality and quantity control - docking VCP/p97 and supporting V-ATPase assembly - to promote degradation rather than exosomal discharge of pro-inflammatory cargo (e.g., angiopoietin-2, cathepsin-D). Its endothelial ablation produces pulmonary and cardiac inflammation, severe sepsis and worsened COVID-19^9^. Endothelial CD82 and CD151 thus constitute a mechanistically antagonistic pair in vasculature, raising the hypothesis that the *balance* between them, probably measurable in blood transcriptomes, could index septic severity and outcome.

A recent transcriptional meta-analysis in sepsis clustered detectable TSPAN genes and identified erythrocyte-related biomarkers (XK, CA1, AHSP, and GYPA) but did not test CD82 or CD151 specifically, did not resolve cell types, and did not examine relationships of tetraspanins to patient survival^10^. We addressed four questions: first, is tetraspanin expression polarized in human sepsis, and does the family move as a coordinated program? Second, does the ratio of this known antagonistic pair (CD82/CD151) predict 28-day sepsis mortality? Third, to what extent are bulk-blood signals driven by changes in cell-type composition versus cell-intrinsic regulation? Finally, do inherited variants that constitutively alter blood tetraspanin transcript abundance also alter risk of developing sepsis?

We find that tetraspanins do not alter in their expression during sepsis as a coordinated family, and that the bulk CD82:CD151 ratio does not correlate with mortality. Instead, sepsis reorganizes tetraspanin expression along two cell-type-specific programs - monocyte CD82 up-regulation and a loss of CD37 across leukocytes - both obscured in bulk whole-blood profiles, and of which only CD37 carries a composition-independent mortality association. Co-expression analysis further places CD82 and CD151 in distinct network modules, consistent with their mechanistic antagonism.

## Methods

### Datasets

Publicly available blood transcriptomic datasets were obtained from the Gene Expression Omnibus (GEO). The bulk meta-analysis comprised eight intensive care unit (ICU)-sampled cohorts (six adult, two pediatric). These were GSE28750, GSE9960, GSE54514, GSE13015, GSE30119, GSE66099, GSE185263 and GSE279448 (241 controls and 550 sepsis patients). All series were screened for cross-series patient overlap, repeated measures, multi-platform duplication and duplicated controls, and every analysis set contains one sample per patient (Supplementary Table S2).

### Dataset sample-specific annotation considerations

GSE185263 enrolled patients from the ICU and emergency department (ED)^11^, and here, only its 82 intensive-care patients (80 with recorded in-hospital mortality) were used, so that every contributing cohort is ICU-sampled like the discovery and external-validation cohorts. Its 266 emergency-department patients, drawn within about two hours of presentation, are reported separately. In GSE30119, a cohort of children with invasive Staphylococcus aureus infection, only the 9 samples flagged sepsis in the deposited sepsis field were counted as sepsis; the remaining 90, predominantly osteomyelitis, abscess and bacteremia without organ dysfunction, were excluded. Survival and co-expression analyses used GSE65682 (Affymetrix U219/GPL13667, whole blood, 802 samples, 28-day mortality available for 468 [103 deaths])^12^. It serves as the discovery cohort throughout; GSE95233, GSE54514 and the GSE185263 intensive-care arm are referred to as the independent ICU cohorts. This dataset was excluded from the case-control meta-analysis because its community-acquired, hospital-acquired and non-pneumonia structure makes a clean sepsis-versus-healthy contrast ambiguous. External mortality validation used GSE95233 (first-day samples, n=51, 17 deaths) and GSE54514 (first-day samples, n=35, 9 deaths). In-hospital mortality in the GSE185263 intensive-care arm (n=80, 20 deaths), which deposits a binary outcome without time-to-event, was modeled by logistic regression (odds ratio [OR] per standard deviation [SD]) and by area under the receiver operating characteristic curve (AUC) against the same random-gene controls; its emergency-department arm (n=265, 32 deaths) was modeled identically and reported alongside it. Single-cell analysis used three 10x Genomics peripheral blood mononuclear cell (PBMC) single-cell RNA-sequencing datasets: GSE167363^13^ (12 samples from 7 donors), GSE279452^27^ (133 samples; the single-cell arm of the study that deposited the bulk series GSE279448, with which it shares 84 donors) and GSE290679^14^ (47 donors: healthy volunteers and ICU patients with and without sepsis). One GSE279452 sample and two GSE290679 samples whose expressed sex contradicted the deposited sex were excluded. Datasets with in vitro stimulation, isolated cell subsets, non-blood tissue, absent healthy controls, or fewer than 10 total samples were excluded.

### Two GSE datasets had substantial annotation problems

First, for GSE54514, the processed values deposited in GEO do not appear to be assigned to the correct samples: Sex, as annotated in the metadata, agrees with sex-specific gene expression at chance levels, serial samples from the same participant differ in expressed sex, and canonical sepsis-induced genes do not differ between its septic and control samples. After rebuilding its expression from the deposited non-normalized raw data, whose columns are BeadChip barcodes that each sample record cites (using limma’s neqc), we find deposited sex then agrees with expression in 99% of samples and sepsis and control samples separated when clustered, showing that the underlying data is intact, but the annotations somehow got mixed up (Supplementary Figure S6). Second was GSE66099, a de-duplicated Day-1 compilation of the unique patients in six earlier Genomics of Pediatric SIRS and Septic Shock Investigators (GPSSSI) series - GSE4607, GSE8121, GSE9692, GSE13904, GSE26378 and GSE26440 - assembled and deposited for that purpose and flagged in GEO as a third-party reanalysis^15^. Those six series contain no Day-1 patients absent from GSE66099, so analyzing them alongside it would count the same 276 children up to three times. We retained GSE66099 and excluded its constituents. Some other studies using these accessions do not appear to have accounted for this overlap – two even used the parent GSE66099 to train a model, which was then validated upon its component studies and, unsurprisingly, both reported very high AUCs^16,17^. The compilation carries fresh sample accessions, so the overlap is not visible by accession matching. It was identified instead from the GEO third-party-reanalysis flag, the series description and the provenance in the supplementary file names.

### Meta-analysis

Probes were collapsed to genes (limma::avereps), log₂-transformed and quantile-normalized. Standardized mean differences (Hedges g) between sepsis and controls were pooled by DerSimonian-Laird random-effects models with Hartung-Knapp-Sidik-Jonkman variance correction (R package meta), with heterogeneity reported as I² and leave-one-cohort-out estimates. CD82 and CD151 were pre-specified from the antagonistic-pair hypothesis; all tetraspanins, including these two, are reported under Benjamini-Hochberg FDR across 33 genes. Adult-only (k=6) and pediatric-only (k=2) sensitivity analyses were run, as were whole-blood-only and PBMC-only strata. A pre-registered family-level test compared the 33 estimable tetraspanins with 1,000 random gene sets matched on platform coverage, expression mean and variance and pooled identically, on three pre-declared metrics: mean |g|, bimodality (bimodality coefficient and Hartigan dip) and cross-cohort sign consistency. A post hoc myeloid/lymphoid lineage bias index - log2(mean expression in monocytes and dendritic cells / T, B and natural killer (NK) cells) from the pseudobulk of each single-cell dataset, defined blind to the meta-analysis - was correlated with pooled g by Spearman with label permutation. In GSE65682, platelet and immature-neutrophil transcript content were each scored as the mean z-score of marker genes (PPBP, PF4, GP9, ITGA2B, TUBB1, GP1BB, SPARC and CLU; MPO, ELANE, CEACAM8, DEFA4, LCN2, LTF and MMP8) and correlated (Pearson) with CD63 among the 685 patients.

### Survival analysis

For GSE65682, Cox proportional-hazards models related per-standard-deviation tetraspanin expression to 28-day mortality. The pre-specified CD82:CD151 ratio was defined as log_2_(CD82) − log_2_(CD151). Because a log-ratio constrains the two genes to equal and opposite coefficients, the pair was additionally tested without that constraint - as an unconstrained bivariate Cox model, with a multiplicative interaction term, as a two-dimensional penalized spline, and by tree-ensemble models - each benchmarked against 1,000 expression- and variance-matched random gene pairs. Hazard ratios (HR), 95% confidence intervals and BH-FDR across testable tetraspanins are reported. Models were re-fitted with the demographic and diagnostic covariates deposited with the cohort (age, sex, admission diagnosis, diabetes). Severity scores were not available. Proportional hazards were assessed by scaled Schoenfeld residuals (cox.zph). For comparison across cohorts, death was related to per-SD CD37 expression by logistic regression in each cohort recording mortality (28-day status in GSE65682), and the three independent ICU cohorts were pooled by inverse-variance meta-analysis, with the Hartung-Knapp p-value also given.

### Single-cell RNA-sequencing

The three datasets were processed identically, one sample at a time (Seurat v5.0.1). GSE167363 and GSE279452 were analyzed from the deposited filtered count matrices; GSE290679 is deposited unfiltered, and its cells were called with the Cell Ranger algorithm (DropletUtils emptyDropsCellRanger, FDR≤0.01). After quality control (200-6,000 detected features, <20% mitochondrial reads; Supplementary Figure S4), doublets were removed with scDblFinder (7.5-11.2% of cells per dataset) and counts were corrected for ambient RNA with DecontX (celda), leaving 1,604,899 cells (55,244 in GSE167363, 861,357 in GSE279452 and 688,298 in GSE290679). Cell types were assigned by mapping each cell to the Azimuth human PBMC reference and grouping its level-2 labels into 14 populations. Reference mapping replaced labeling by marker-module scores (AddModuleScore), under which 36% of the cells called NK in GSE279452 carried a productive T-cell receptor, against 2% after mapping. With the mapped labels 98% of receptor-bearing T cells were labeled as T cells, and in GSE290679 the labels agreed with those deposited by its authors for 94% of cells. Because the unit of biological replication is the donor rather than the cell, corrected counts were aggregated to donor x cell-type pseudobulk values (log2 counts-per-million), excluding donor x cell-type units with fewer than 10 cells. Differences between groups are reported as the difference in mean log2 counts-per-million with Hedges g and its 95% confidence interval; p-values are from Welch t-tests, with Benjamini-Hochberg correction across the comparisons displayed for each dataset. For Figure 4A the three datasets were pooled in a linear model of donor-level log2 counts-per-million on sepsis status with dataset as a covariate, each donor counted once, with Benjamini-Hochberg correction across the 108 comparisons shown; a comparison is marked only if it also has the same direction in every dataset. At ICU admission (T0) GSE167363 comprises two healthy donors, three survivors and two non-survivors. Comparisons within GSE167363 are therefore reported descriptively rather than as powered hypothesis tests, and their stability was assessed by leave-one-donor-out. GSE279452 contributed 7 healthy and 92 septic adults and 8 non-septic adult ICU patients; its 25 children, four of them healthy, were not contrasted. GSE290679 contributed 9 healthy donors and 19 septic and 17 non-septic ICU patients. Outcome was deposited only with GSE167363. Single-cell analyses used R 4.3.2 with scDblFinder 1.16.0, celda 1.18.2, Azimuth 0.5.1 and DropletUtils 1.22.0.

### Co-expression network

Weighted gene co-expression network analysis (WGCNA) was applied to the 3,000 most variable genes in 685 intensive-care patients (GSE65682) using a signed network, Pearson correlation, and soft-thresholding power 14. GSE65682 deposits 84 controls (42 healthy volunteers, 42 non-infected surgical intensive-care controls) and 718 intensive-care patients; controls and the 33 patients labeled non-pneumonia were excluded, leaving 685. Modules were related to 28-day survival, time-to-event and tetraspanin expression. Intramodular hub genes and Gene Ontology enrichment (clusterProfiler) were computed.

### Deconvolution and composition-adjusted survival

Leukocyte fractions were estimated with EPIC (immunedeconv) using the circulating-blood reference (BRef; tumor=FALSE), which returns B cells, CD4+ and CD8+ T cells, monocytes, neutrophils, NK cells and an uncharacterized fraction. Per-cell-type log2 fold-changes were pooled by random-effects meta-analysis across the five independent cohorts with sufficient healthy controls for deconvolution. Because EPIC fractions are compositional and constrained to sum to a constant, they index relative rather than absolute changes, and covariate adjustment for them does not resolve that constraint. In GSE65682, Cox models for tetraspanin expression were re-fitted with all estimated cell fractions as covariates to distinguish cell-intrinsic from cell type-compositional effects. Analyses used R 4.3.2 and 4.5.1, code available at https://gitlab.com/jdwren/sepsis_tetraspanin2.

### Machine-learning models and null benchmarking

To test whether the tetraspanin panel as a whole, rather than any single gene, carries prognostic information, we fitted a penalized (elastic-net) logistic regression predicting 28-day mortality in GSE65682 from the 23 tetraspanins present on the array, chosen by array coverage rather than by outcome, with the penalty mixing parameter and strength tuned by cross-validation. Discrimination within the cohort was estimated by repeated nested cross-validation (ten subsets, ten repeats, deaths balanced), scoring only held-out patients, and summarized as the AUC (where 0.5 is chance) with 95% confidence intervals. We then tested the final model on two independent cohorts (GSE95233 and GSE54514, first-day samples) using the tetraspanins shared by both platforms. Because intensities are not comparable across platforms, values were converted to within-sample ranks, with quantile normalization as a sensitivity check. Because an external AUC is interpretable only against what arbitrary genes achieve on the same comparison, every external result was compared against three controls generated by rerunning the analysis with only the input genes changed. These were all 10,009 shared genes, 1,000 random 23-gene panels matched on mean expression and variability, and the distribution of single-gene AUCs across all shared genes. Random forests and gradient boosting were run identically, and any result specific to one model was required to replicate on a second external cohort (GSE54514) before being credited. Bootstrap resampling (1,000 resamples, lasso penalty) assessed whether any individual tetraspanin was consistently retained. Given few external events, we treat rankings against these controls as more reliable than the absolute AUCs.

### Mendelian randomization

To test whether inherited tetraspanin transcript abundance causally affects sepsis susceptibility, we performed summary-data Mendelian randomization (SMR with the HEIDI heterogeneity test^18^) together with Bayesian colocalization^19^ (coloc.abf). This is a single-instrument cis design rather than a conventional multi-instrument analysis. cis-expression quantitative trait locus (cis-eQTL) instruments were drawn from eQTLGen whole blood^20^ (n=31,684, primary), with GTEx^21^ v8 whole blood (n=670) as a sensitivity exposure. The outcome was the FinnGen^22^ R12 sepsis endpoint (AB1_OTHER_SEPSIS, 17,133 cases and 439,048 controls), using 1000 Genomes Phase 3 Europeans as the linkage-disequilibrium reference. Of the 33 canonical tetraspanins, 24 carried a genome-wide-significant whole-blood cis-eQTL instrument (instrument p≤5×10⁻⁸) and were testable. The remaining nine lacked a cis probe in eQTLGen and were not instrumentable (Supplementary Table S1). SMR p-values were FDR-controlled (Benjamini-Hochberg) across the 24 testable genes, HEIDI p<0.01 was taken to indicate linkage or pleiotropy rather than a shared causal variant, and colocalization required a posterior probability of a shared causal variant (PP.H4) ≥0.75. Because instrument strength bounds bias rather than power, the minimum effect detectable with 80% power at α=0.05 was computed directly from each gene’s SMR standard error. To confirm that the pipeline detects a true effect when one exists, we ran a tissue-matched positive control - the blood/monocyte-mediated locus LIPA against the FinnGen R12 coronary-heart-disease endpoint. Analyses used SMR v1.4.1.

### Experimental single-cell validation (mouse)

As an orthogonal, cross-species test of the CD82 and CD151 axes, we analyzed single-cell RNA-sequencing of mouse lung following lipopolysaccharide (LPS) challenge versus control. Cells were processed in Seurat and annotated to nine pulmonary cell populations, and expression of four tetraspanins (Cd37, Cd53, Cd82 and Cd151) was compared between LPS and control within each cell type (Wilcoxon rank-sum test, fraction of expressing cells and log2 fold-change reported). To increase cell yield, four mice were pooled per sample and two pooled samples were analyzed per group. Because individual-mouse replicates are therefore not resolvable, between-condition comparisons are interpreted descriptively rather than as powered hypothesis tests.

## Results

### Tetraspanin expression changes in sepsis track myeloid versus lymphoid lineage composition

Across eight ICU-sampled cohorts, several tetraspanins showed large, directionally consistent expression changes in sepsis (Figure 1). CD63 (Hedges g +2.07), CD82 (+1.39), CD151 (+0.89), ROM1 (+0.66), TSPAN9 (+0.48) and UPK1A (+0.35) were up-regulated, whereas CD81 (−1.72), TSPAN3 (−1.51), TSPAN13 (−1.20), TSPAN32 (−0.88), TSPAN18 (−1.09), TSPAN4 (−0.62) and TSPAN31 (−0.67) were down-regulated. These 13 genes were significant after FDR correction across the 33-gene family (FDR<0.05), with the same direction in 93 of the 97 gene-by-cohort estimates (Supplementary Figure S1). The family as a whole, however, was not polarized: against 1,000 matched random gene sets, the 33 tetraspanins were no more perturbed (mean |g| 0.588, 26th percentile of the null), no more split into up and down blocks (6th percentile), and no more reproducible in direction across cohorts (0.804, 43rd percentile) than matched random genes. Adult-only and pediatric-only analyses were concordant in direction for all 13 genes, as were whole-blood-only and PBMC-only analyses. Both CD82 and CD151 rose in sepsis and both were significant across the 33-gene family (FDR=0.003 and 0.005, respectively), with the larger effect on CD82 (g=+1.39 vs +0.89), shifting the net balance in sepsis toward CD82. CD63 showed the largest pooled effect (g=+2.07, 95% CI +1.15 to +2.98, FDR=0.016).

**Figure 1.**
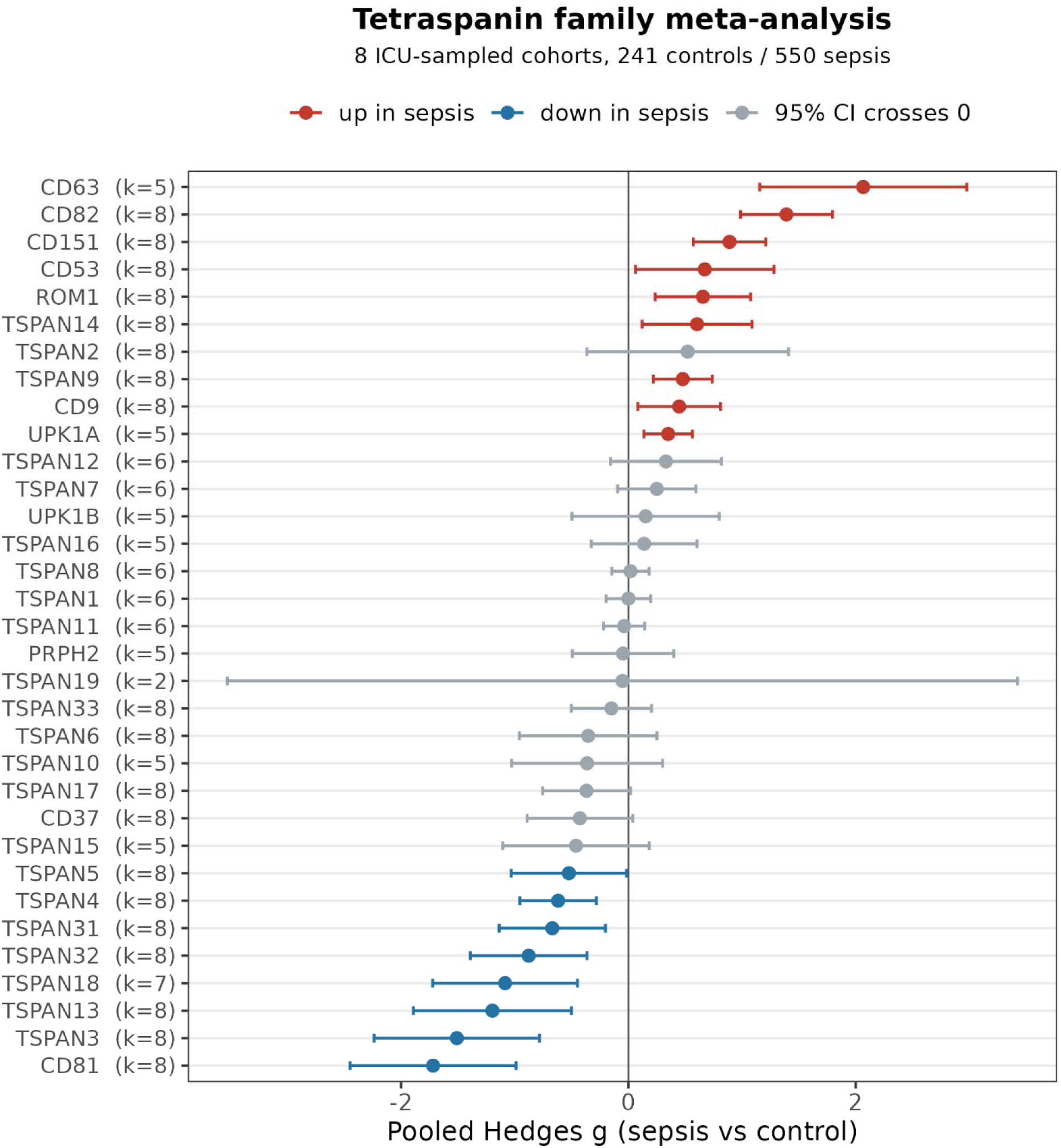
Pooled standardized mean differences (Hedges *g*, random-effects) for tetraspanin genes in sepsis versus controls across the eight ICU-sampled cohorts (241 control and 550 sepsis patients; per-gene analyzed N varies with platform coverage). Points are pooled estimates, bars are 95% confidence intervals. Red, up-regulated in sepsis; blue, down-regulated; gray, the 95% confidence interval crosses zero. 17 genes have an interval excluding zero, of which 13 also reach FDR<0.05 across the family. Both CD82 and CD151 increase, with a larger CD82 effect.

Two properties distinguished the tetraspanins that changed. First, they were the ones abundant enough in leukocytes to measure. Genes with |*g*|≥0.5 averaged 4.55, 5.29 and 5.33 log2 counts-per-million in the three single-cell datasets (GSE167363, GSE279452 and GSE290679, in that order here and below) against 2.44, 2.57 and 2.62 for the rest (p=0.008, 0.002 and 0.001), while much of the family did not have detectable expression in these datasets. Second, among the abundant genes, the direction of change was associated with lineage-biased expression. A myeloid/lymphoid lineage bias index correlated with the pooled effect (Spearman ρ=0.53, 0.41 and 0.22 among high-effect genes, ρ=0.43, 0.42 and 0.33 family-wide, permutation p=0.012, 0.014 and 0.050; Supplementary Figure S7). Tetraspanins whose expression is biased toward myeloid cells among circulating leukocytes rose (CD63, CD151), and those biased toward lymphocytes fell (TSPAN32, TSPAN18), mirroring the myeloid expansion and lymphoid depletion of sepsis. CD82 was the exception: its index was positive only in GSE167363 (+2.08, against −0.82 and −1.10 in the two larger datasets), and its increase in blood instead followed from induction within monocytes (below). Much of what bulk blood reports for tetraspanins is therefore shifting leukocyte composition rather than a coordinated membrane-domain program.

### The CD37 mortality association reproduces in independent ICU cohorts

Contrary to our *a priori* hypothesis, the bulk CD82:CD151 ratio did not predict 28-day mortality (HR=1.09 per SD, 95% CI=0.90-1.33, p=0.37), nor did CD82 (HR=1.09 per SD, FDR=0.63) or CD151 alone in the full cohort (Figure 2A). Relaxing the log-ratio’s equal-and-opposite constraint (unconstrained bivariate Cox, interaction, penalized-spline and tree-ensemble models) was equally uninformative, and the pair ranked at the 12th percentile of 1,000 expression-matched random gene pairs. The strongest survival-associated tetraspanin was CD37, with lower expression associated with higher mortality (HR=0.72 per SD, 95% CI=0.60-0.85, FDR=0.002; Figure 2A). Adjustment for the demographic and diagnostic covariates deposited with the cohort left the CD37 association essentially unchanged (HR=0.73, p<0.001). Because bulk blood signals can reflect changing leukocyte proportions, we estimated cell fractions and re-fitted survival models with composition as covariates. Sepsis was accompanied by a directional expansion of monocytes and a corresponding reduction in lymphocyte fractions, with NK-cell, B-cell and CD8+ T-cell depletion reaching FDR<0.05 across the five cohorts available for deconvolution (Figure 5). The CD37 hazard ratio was unchanged after composition adjustment (0.72 => 0.70; p=0.007, FDR=0.029 across the four adjusted models), consistent with a composition-independent association, whereas the weaker CD81 association attenuated to non-significance (0.79 => 0.85), consistent with a largely compositional origin (Figure 2B).

**Figure 2.**
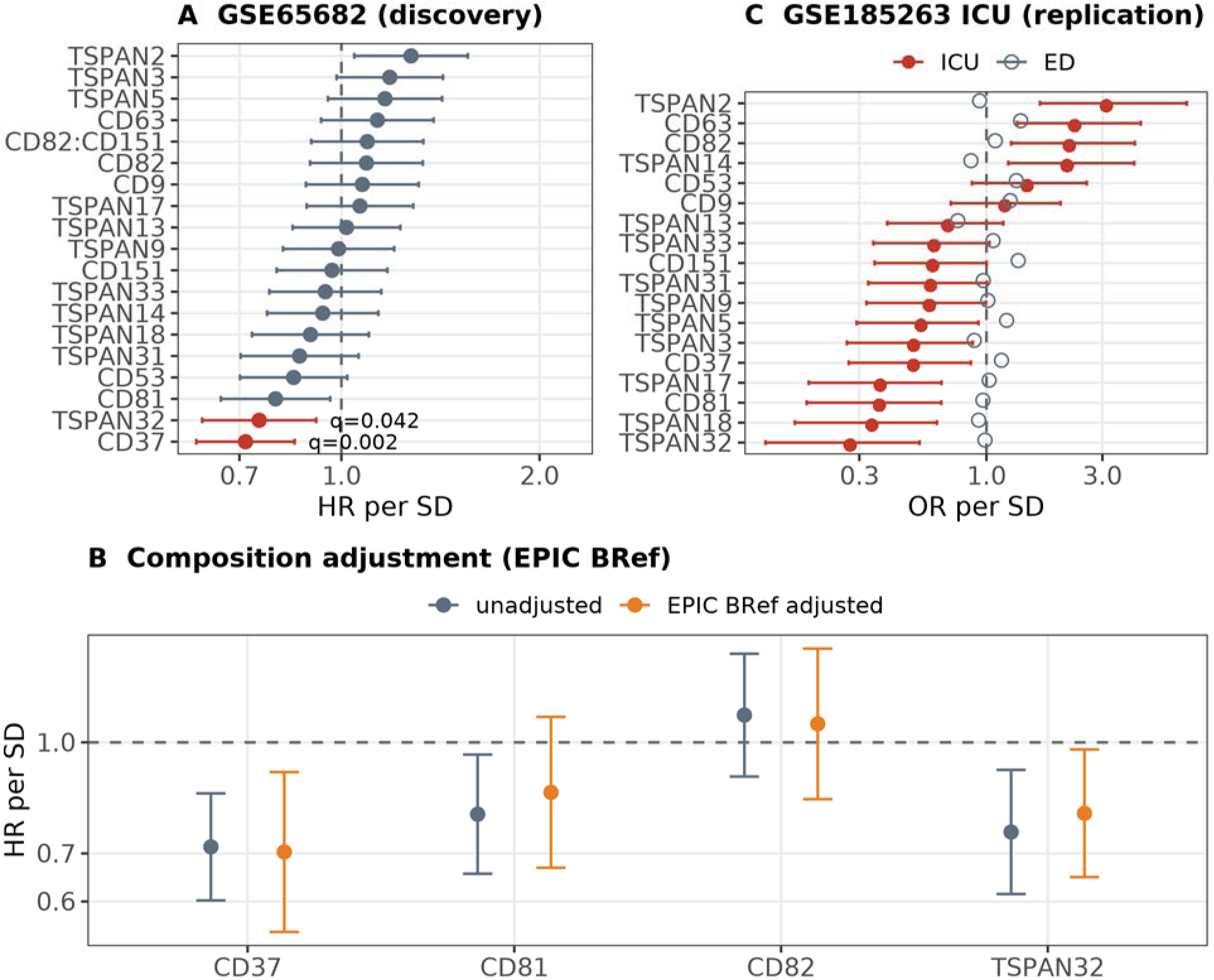
(A) Cox proportional-hazards ratios for 28-day mortality per SD of tetraspanin expression (GSE65682, n=468, 103 deaths), for the 18 tetraspanins detected above array background; points are HR, bars 95% CI, q = FDR across those 18 genes, the dashed line marks HR=1. Red marks q<0.05. (B) Hazard ratios before (gray) and after (orange) adjustment for EPIC-estimated leukocyte fractions, using the circulating-blood reference (BRef), which includes the neutrophil and monocyte fractions. (C) ORs per SD for in-hospital mortality in the GSE185263 intensive-care arm (n=80, 20 deaths; filled red), the independent replication cohort, with its emergency-department arm (n=265, 32 deaths; open gray) shown alongside. This cohort deposits a binary outcome without time-to-event, so an OR rather than a hazard ratio is shown.

To test whether the tetraspanin panel, as a whole, contains prognostic information, we trained a regularized (elastic-net) classifier for 28-day mortality. Cross-validation selected a pure ridge penalty (alpha=0), so no sparse subset of the panel was preferred. The panel achieved only modest internal discrimination (repeated cross-validation AUC=0.61, 95% CI=0.55-0.67). Discrimination did not generalize to an independent cohort (GSE95233; external AUC=0.48, 95% CI=0.28-0.68), and this failure was robust to the cross-platform normalization method. A gradient-boosted model reached an external AUC of 0.775 on this cohort, but the signal was additive, distributed across the family rather than concentrated in any gene, and did not replicate on a second external cohort (GSE54514, AUC=0.509, p=0.821). By comparison, a whole-transcriptome model retained transferable discrimination (AUC=0.70, 95% CI=0.54-0.86) and expression-matched random 23-gene panels achieved a median external AUC of 0.62, so the pipeline carries transferable mortality signal that the tetraspanin panel does not. Tested individually, the CD37 association kept its direction in GSE95233 (AUC=0.645, 95% CI=0.489-0.801) and in GSE54514 (AUC=0.70, 9 deaths). In neither cohort, however, did CD37 differ significantly from the single-gene distribution of the whole transcriptome (85th and 93rd percentiles; empirical p=0.15 and p=0.075), and cross-cohort single-gene replication between these two external sets was itself absent (67 genes ranked in the top decile of both, against 88 expected by chance). In the GSE185263 intensive-care arm (80 patients, 20 in-hospital deaths), higher CD37 expression was associated with lower mortality (OR=0.50 per SD, 95% CI=0.27-0.86, p=0.018; AUC=0.670, 95% CI=0.517-0.823), reproducing the direction and approximate magnitude of the discovery-cohort effect (Figure 2C). This replication was not specific to CD37: 10 of the 18 detected tetraspanins reached FDR<0.05 among the same 20 deaths, several more strongly than CD37 (FDR=0.035), and CD37 sat at the 80th percentile of the transcriptome-wide AUC distribution. Higher CD82 expression was associated with higher mortality in the same arm (OR=2.19, FDR=0.020). In the emergency-department arm of the same cohort neither was associated with death (CD37 OR=1.15, p=0.46; CD82 OR=1.09, p=0.66). Pooled across the three independent ICU cohorts (166 patients, 46 deaths), the CD37 OR was 0.56 per SD (95% CI=0.38-0.81, p=0.002; Hartung-Knapp p=0.018; I²=0%; Figure 3). The CD37 mortality association therefore reproduces in intensive-care sepsis, and the tetraspanin family carries no multivariate mortality signal beyond it.

**Figure 3.**
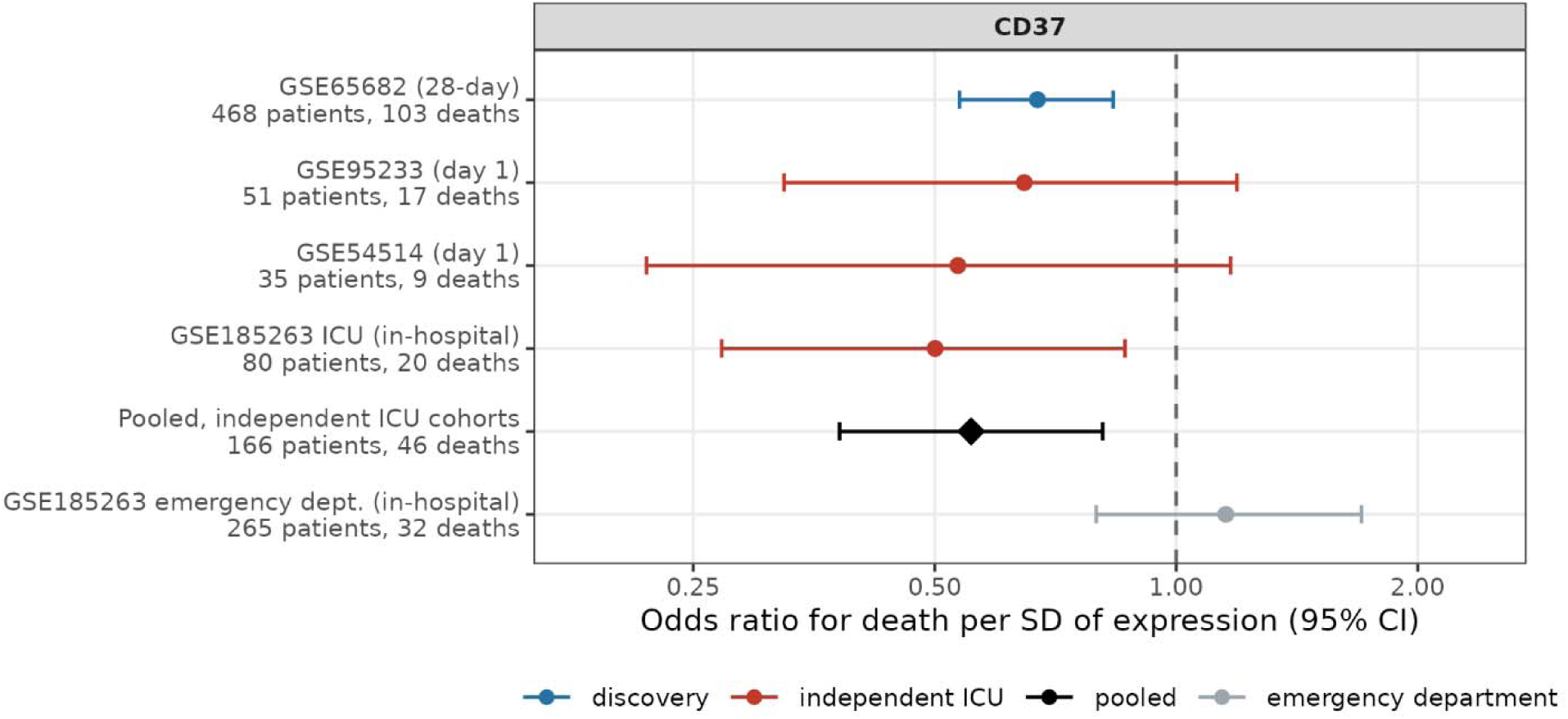
OR for death per SD of CD37 expression (logistic regression - points are OR, bars 95% CI) in each cohort recording mortality: the discovery cohort GSE65682 (blue; 28-day status), the three independent ICU cohorts (red; GSE95233 and GSE54514, first-day samples; GSE185263 intensive-care arm, in-hospital mortality), their inverse-variance pooled estimate (black diamond; OR=0.56, 95% CI=0.38-0.81, I²=0%, p=0.002, Hartung-Knapp p=0.018) and the GSE185263 emergency-department arm (gray), which is not pooled. The dashed line marks OR=1.

### Single-cell analysis resolves two cell-type-specific tetraspanin programs

Donor-level analysis of 1.6 million PBMCs from three datasets (Supplementary Figure S2) showed that bulk signals are superpositions of opposing cell-type-specific programs (Figure 4A). CD82 up-regulation in sepsis was reproduced in CD14 monocytes in all three datasets (+0.63, +1.74 and +1.56 log2 in GSE167363, GSE279452 and GSE290679; 95% CI excluding zero in the two larger) and accompanied by down-regulation in B cells (−1.32, −0.50, −0.75) and CD4+ T cells (−1.90, −0.48, −0.63), so that the whole-blood CD82 increase reflects both monocyte induction and shifting cell proportions. NK cells were the exception among lymphocytes: CD82 was higher in sepsis in all three datasets (+0.78, +1.62, +0.57; 95% CI excluding zero in GSE279452 only). CD151 did not change reproducibly: it rose in 7 of 8 populations in GSE167363 (CD14 monocytes +0.81), did not differ from zero in GSE279452 (CD14 monocytes +0.31, B cells +0.01) and fell in 9 of 12 in GSE290679 (CD14 monocytes −0.47, B cells −0.62; both intervals excluding zero), so the whole-blood CD151 increase is not explained by induction within mononuclear cells. Only in platelets was it higher in all three (+0.47, +0.59, +0.28). CD63, the largest effect in the bulk meta-analysis, was induced within monocytes in all three datasets (pooled +1.08 log2 in CD14 and +1.65 in CD16 monocytes; Figure 4A), and CD14 monocytes carried 33-61% of the CD63 transcripts of septic PBMCs. It was also higher within platelets (+0.62), but platelets held under 4% of those transcripts, and in whole blood CD63 tracked the granule-protein transcripts of immature neutrophils (r=0.39 among 685 septic patients) rather than platelet transcripts (r=0.02). In contrast, CD37 was almost uniformly down-regulated across all leukocyte types, most in CD14 monocytes (−2.36, −0.91, −0.82), explaining its low variance in bulk data. The CD82 and CD37 directions matched the pooled estimate in the eight-cohort meta-analysis, a far larger sample for the sepsis-versus-healthy contrast than the 18 healthy donors profiled here. CD37 itself was not significant after FDR correction in the bulk meta-analysis, so its pan-leukocyte decrease rests on these single-cell comparisons.

**Figure 4.**
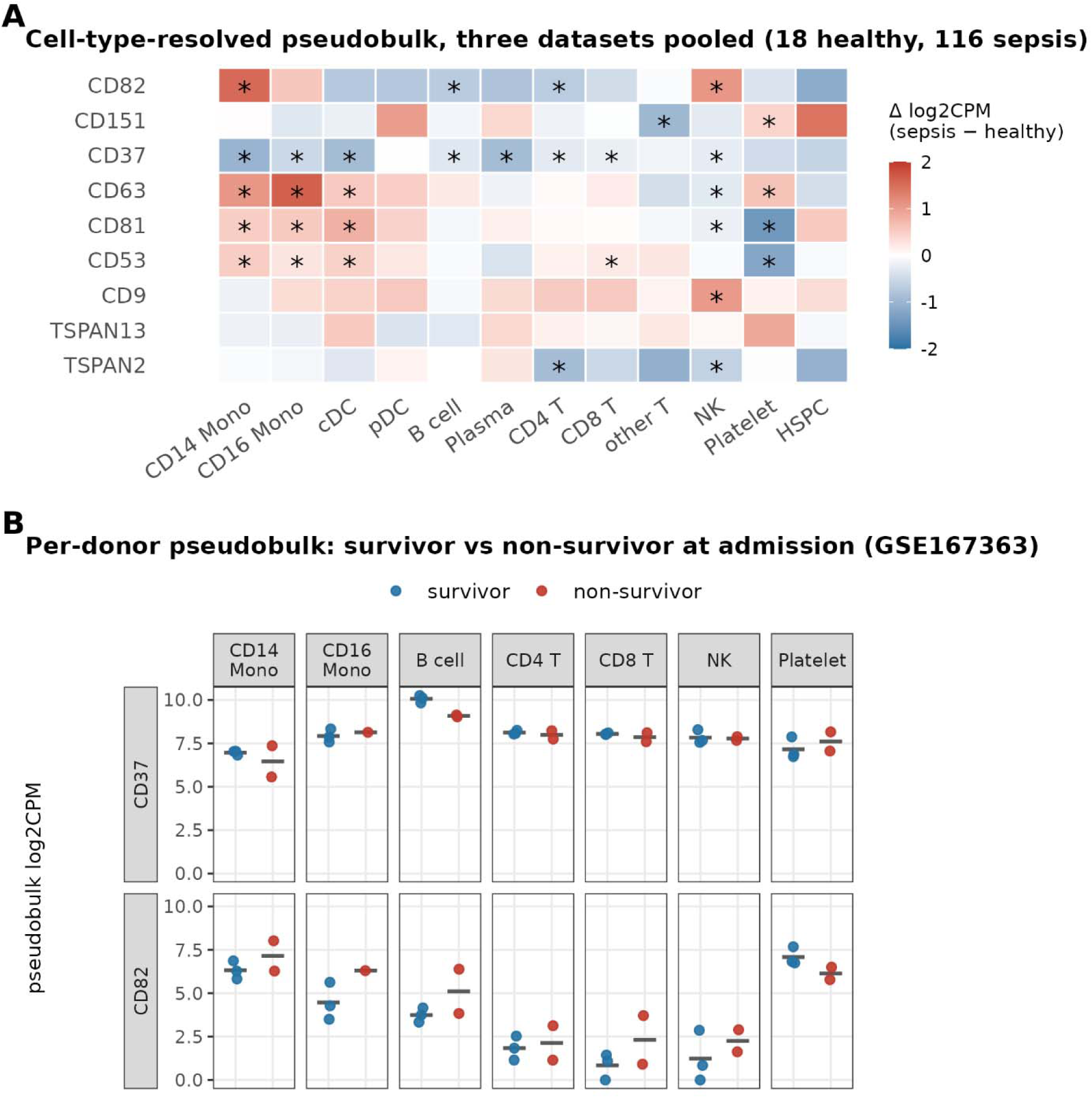
(**A**) Cell-type-resolved differential expression (donor × cell-type pseudobulk log2 fold-change, sepsis vs. healthy) for focal tetraspanins across PBMC populations pooled across the three datasets (18 healthy and 116 septic donors; the difference within dataset, each donor counted once). Asterisks mark comparisons at FDR<0.05 that have the same direction in every dataset. CD82 rises in CD14 monocytes and falls in B and CD4+ T cells in all three datasets, CD37 falls across leukocytes, and CD151 shows no pooled change in monocytes, B cells or CD4+ and CD8+ T cells, where the datasets disagree (Supplementary Figure S8). (**B**) Per-donor pseudobulk values in non-survivors (2) vs survivors (3) at admission in GSE167363, each point one donor. Comparisons are underpowered and reported descriptively. Panel A displays 108 gene by cell-type comparisons, of which 32 are marked, among them CD82 and CD37 in CD14 monocytes.

**Figure 5.**
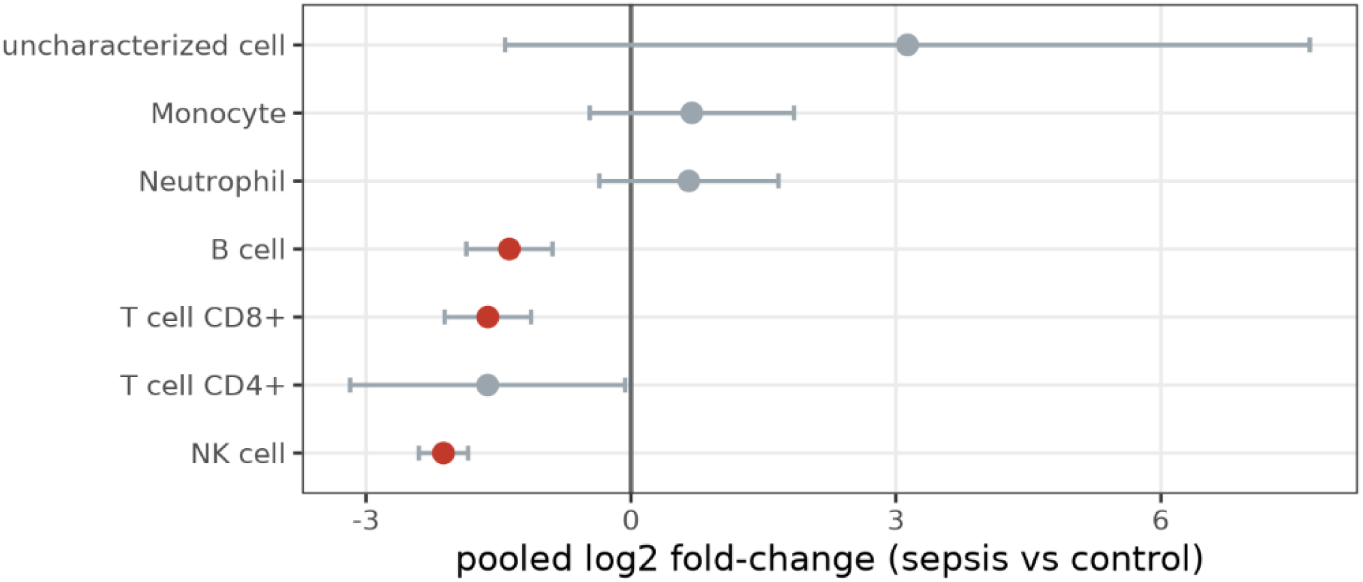
Random-effects pooled leukocyte-fraction changes in sepsis (EPIC) across the five whole-blood cohorts with sufficient healthy controls for deconvolution, showing depletion of NK cells (pooled log2 fold-change −2.12, FDR<0.001), B cells (−1.38, FDR=0.003) and CD8+ T cells (−1.62, FDR=0.003), with directional monocyte expansion. Fractions are compositional and index relative rather than absolute change.

The two larger datasets also profiled ICU patients without sepsis, who fell between healthy donors and septic patients. In CD14 monocytes CD82 was already higher in non-septic ICU patients than in healthy donors (+1.30 in GSE279452, +0.88 in GSE290679; both intervals excluding zero) and higher again in sepsis (+0.44 and +0.68; interval excluding zero in GSE290679 only), and CD37 fell in two steps (−0.52 and −0.39, then a further −0.39 and −0.43; all four intervals excluding zero), and CD63 was likewise already raised (+1.30 and +0.58; both intervals excluding zero). Part of each program is therefore a response that starts in critical illness.

Outcome was deposited only with GSE167363, which at admission comprised three survivors and two non-survivors (Figure 4B). CD37 was lower in non-survivor B cells (−0.98 log2), in every donor and in every leave-one-donor-out fold, concordant in direction with the composition-independent CD37 mortality association measured in the much larger bulk cohort. Plasmablasts (118 cells, 0.2% of the dataset after reference mapping) were too few to compare. CD82 was higher on average in non-survivor CD14 monocytes (+0.83 log2), although the two groups overlapped and the direction did not hold in every leave-one-donor-out fold, and CD82 carried no mortality association in the discovery cohort, GSE95233 or GSE54514. These single-cell data are consistent with a lymphoid origin for the bulk CD37 mortality association but, with five donors, cannot establish it.

### CD82 and CD151 anchor distinct co-expression modules

To test whether the pair is separated at the network level, we performed WGCNA in 685 intensive-care patients. CD82 and CD151 were hub genes of *different* modules - CD82 in a myeloid module (module-gene r=0.752) and CD151 in a separate module (r=0.633) - while CD81 anchored a separate lymphocyte-associated module (r=0.793) (Supplementary Figure S3). The CD82 module is annotated to myeloid activation, inflammatory signaling and neutrophil granule genes (module-membership annotation rather than formal enrichment testing) and contains SEMA4A, MAPK14 (p38α), ALOX5 (5-lipoxygenase) and NLRC4, four myeloid effectors whose activity depends on membrane localization rather than abundance alone. Their co-clustering with CD82 is consistent with a module organized around membrane-proximal myeloid activation, though module membership is correlative and does not establish that CD82 organizes these proteins. CD82 and CD151 showed no transcriptional coupling beyond the background correlation shared by arbitrary expression- and variance-matched gene pairs in these patients (r=+0.15, 56th percentile of 2,000 matched random pairs, p=0.88).

### Myeloid CD82 induction and endothelial CD151 loss are recapitulated in an LPS-challenged mouse lung

To test whether the two membrane axes generalize beyond human circulating blood, we examined an independent single-cell RNA-sequencing dataset of mouse lung following LPS challenge, profiling Cd37, Cd53, Cd82 and Cd151 across nine pulmonary cell populations (Table 1, Supplementary Figure S5). Because this model differs from human sepsis in species, tissue and stimulus, and cells were pooled, comparisons are descriptive. Two observations mirrored the human findings. First, Cd82 was induced in macrophages (11% to 56% of cells expressing after LPS) but unchanged in NK cells and neutrophils, paralleling the monocyte CD82 up-regulation in human sepsis. Second, Cd151 was essentially confined to endothelial cells - a cell type absent from peripheral-blood profiling - where the fraction of expressing cells fell from 51% to 29% after LPS, consistent with loss of the vascular-protective tetraspanin during endothelial injury.

**Table 1.** Tetraspanin detection in LPS-challenged mouse lung (single-cell RNA-sequencing). Percent of cells with detectable transcript in control vs LPS-challenged mouse lung. Highlighted rows are the two CD82/CD151 axis-relevant observations. Because cells were pooled (four mice per pool, two pools per group), values are descriptive rather than a powered statistical test.

| Gene | Cell type | Ctrl → LPS | log<sub>2</sub>FC | Pattern |
| --- | --- | --- | --- | --- |
| **Cd82** | **Macrophage** | **10.7% → 56.2%** | **+0.45** | **Induced by LPS (5.3-fold more cells)** |
| Cd82 | NK cell | 43.1% → 44.7% | -0.02 | Expressed, unchanged |
| Cd82 | Neutrophil | 38.4% → 34.7% | -0.10 | Expressed, unchanged |
| **Cd151** | **Endothelial cell** | **50.8% → 28.9%** | **-0.34** | **Reduced by LPS** |
| Cd151 | Other 8 types | all <5% | - | Endothelial-restricted |
| Cd37 | B cell | 75% → 80% | +0.02 | B-cell-enriched, unchanged |
| Cd37 | Other leukocytes | 8% – 47% | ~0 | Broadly expressed, unchanged |
| Cd53 | Most leukocytes | 60% – 98% | ~0 | Pan-leukocyte, unchanged |

The remaining two tetraspanins were unchanged. Cd37 was most highly expressed in B cells, as in humans, but was not altered by acute LPS in any cell type, consistent with the human CD37 change being a feature of established sepsis rather than acute endotoxin exposure. Cd53, included as a leukocyte-restricted comparator, was broadly expressed across immune populations and unchanged. In line with the mouse result, CD53 did not reach family-wide significance in human septic blood (meta g=+0.67, FDR=0.07) and was not associated with mortality (Cox HR=0.85 per SD, p=0.081). These data place Cd82 induction and Cd151 loss in different cell types, consistent with the separation seen in human blood, but they do not test the CD82/CD151 antagonism itself, which has been established functionally in endothelium rather than in leukocytes. Leukocyte Cd151 was near-absent in mouse lung, whereas CD151 was expressed across human PBMC populations without a reproducible change in sepsis, a species or tissue difference that cell-resolved human data will need to resolve.

### Constitutive tetraspanin expression shows no causal effect on sepsis susceptibility

Finally, to distinguish disease-reactive changes from germline causation, we asked whether inherited variants that set constitutive blood tetraspanin expression alter sepsis risk. Among the 24 instrumentable tetraspanins, none showed a significant causal effect on sepsis after FDR correction (all FDR>0.19; smallest raw p, TSPAN13 p=0.009 and CD63 p=0.016), and none colocalized with the sepsis association signal (all PP.H1-dominant - the eQTL is present but there is no sepsis signal at the locus; maximum PP.H4=0.14, TSPAN13). The GTEx whole-blood sensitivity analysis agreed, and UPK1A - absent from eQTLGen but instrumentable in GTEx (F=114) - was likewise null (PP.H4=0.005). Instruments were strong (F=81-10,840; median=518), arguing against weak-instrument bias. Power computed from each gene’s standard error gave 80% power to detect an OR of 1.11 per SD of whole-blood CD37 transcript (median 1.19 across the 24 genes). The tissue-matched positive control (LIPA and coronary heart disease) returned a significant causal estimate (SMR p=2.6×10⁻¹⁰) with strong colocalization (PP.H4=0.99). Genetically proxied baseline whole-blood tetraspanin expression therefore provides no evidence of an effect on sepsis susceptibility, within the effect sizes these instruments can resolve (Supplementary Table S1).

## Discussion

Integrating eight ICU-sampled cohorts with single-cell, network and genetic analyses, we find that tetraspanin expression is extensively reorganized in human sepsis - thirteen of the 33 family members change at FDR<0.05 - and that this reorganization is best understood at the level of cell types rather than as a single bulk ratio. Three conclusions stand out.

First, CD82 and CD151 both rise in septic blood, with CD82’s effect roughly one and a half times as large, yet the bulk ratio does not predict 28-day mortality and the pair performed no better than expression-matched random gene pairs. Paired tetraspanin subsets with opposing roles have been observed before in cancer, where CD9 and CD82 suppress metastasis while CD151 and TSPAN8 promote it^23,24^, CD82 and CD151 being that literature’s canonical antagonistic pair. In vascular inflammation the polarity is reversed - CD82 is protective in metastasis but pro-leakage in endothelium - yet the antagonism persists. The TSPANs with the largest changes segregate along myeloid versus lymphoid lineages, so much of what a bulk panel reports is the leukocyte composition shift of sepsis rather than a coordinated membrane program. This does not contradict the axis framing: opposing-role tetraspanin pairs are defined by function within a cell, established by knockdown and overexpression^23^, not by coordinated transcription across a tissue. Three features of our data explain why the bulk ratio fails. (i) The two genes are not transcriptionally coupled in blood: their correlation across these patients is +0.15, the 56th percentile of expression- and variance-matched random pairs, so they are no more related to one another than two arbitrary genes. (ii) The single-cell analysis shows that CD82 induction is concentrated in monocytes while CD82 falls in B and CD4+ T cells, whereas CD151 shows no reproducible change within mononuclear cells, so the two arms have different cell-type architectures and a whole-blood ratio averages over cell-types with differing biology and shifting proportions. (iii) The mouse data place Cd82 and Cd151 in different cell types: Cd82 rises in macrophages while Cd151 falls in endothelium. The antagonistic axis defined in endothelium and isolated cells^7,9^ is therefore not detectable within unfractionated blood - a caution for biomarker programs built on bulk tetraspanin ratios, and an argument for cell- and tissue-resolved measurement rather than evidence against the axis itself.

Second, cell resolution separates two cell-type-specific programs, only one of which, CD37 loss, correlates consistently with outcome. CD82 induction in monocytes was reproduced in three single-cell datasets and is consistent with a pro-inflammatory role, although the pro-leakage function of CD82 and its coupling to accessible-cholesterol/Rho-GTPase signaling were established in endothelial cells^7^, and whether monocyte CD82 has comparable effects is untested. Its co-expression with innate pattern-recognition genes is correlative. It is a robust marker of the septic state, reproduced in mouse lung macrophages. In both datasets that included non-septic ICU patients their monocyte CD82 was already raised, so the marker is not specific to infection. It is not, however, a consistent mortality signal: CD82 was non-significant in the discovery cohort before and after composition adjustment and in the two smaller external cohorts, and although it was associated with death in the GSE185263 intensive-care arm (OR=2.19 per SD, FDR=0.020), 10 of 18 tetraspanins reached FDR<0.05 in that arm. The single-cell survivor comparison rests on too few donors to establish one.

By contrast, loss of CD37 across leukocytes - lower still in non-survivor B cells - was the strongest survival-associated signal within the discovery cohort, persisting after adjustment for the deposited clinical covariates and for leukocyte composition including the neutrophil and monocyte fractions. CD37 loss is consistent with the humoral component of sepsis-induced immunosuppression and with the B-cell depletion estimated by deconvolution. These cell-type-level shifts are consistent with a recent single-cell multi-omic atlas of 281 adult and pediatric sepsis patients and controls, which reported lymphocyte and natural-killer-cell depletion alongside monocyte expansion^25^, and a pediatric single-cell study identified B cells as the most depleted leukocyte population^26^.

Prior studies have established that CD37 is restricted to the immune system, expressed most highly on mature B cells, and organizes partner proteins into tetraspanin-enriched membrane domains^27^. Two of its partnerships converge on IL-6: CD37 stabilizes the IL-6 receptor complex with suppressor of cytokine signaling 3 (SOCS3) so that feedback inhibition can act, and its loss leaves IL-6/AKT/STAT3 signaling constitutively active^28^. Separately, CD37 associates with the pattern-recognition receptor dectin-1, and CD37-deficient macrophages over-produce IL-6 in response to dectin-1 ligands^29^. Because circulating IL-6 predicts sepsis mortality^30^, CD37 loss that disinhibits IL-6 signaling is a plausible mechanism for the association. Cd37^-/-^ mice mount poor T-cell-dependent IgG responses^31^, show impaired dendritic cell migration and T-cell priming^32^, and recruit neutrophils poorly, because CD37 supports β2-integrin-dependent adhesion and cytoskeletal function^33^, phenotypes that fit the immunosuppressed phase of sepsis. Thus, the association between CD37 loss and sepsis mortality is consistent with established mechanisms. In diffuse large B-cell lymphoma, CD37 loss correlates with IL-6 pathway activation and worse survival^28^ and improves risk prediction independently of the International Prognostic Index^34^. In sepsis the association reproduces but is not specific to CD37. Higher CD37 expression is associated with lower mortality in the GSE185263 intensive-care arm (OR=0.50 per SD, p=0.018), and in the same direction in the two smaller ICU cohorts, yet 10 of the 18 detected tetraspanins reach FDR<0.05 among the same 20 deaths and CD37 ranks below TSPAN18, TSPAN17, CD81 and several others within that arm. Single-gene mortality association in intensive-care sepsis blood is diffuse rather than gene-specific, and a gene that separates survivors from non-survivors there is not thereby unusual.

Third, the co-expression network places CD82 and CD151 in distinct modules. CD82 is strongly co-expressed with the membrane-microdomain organizer flotillin-1 (r≈0.70 in both septic patients and healthy controls, so this is a constitutive relationship rather than a sepsis-induced one). Tetraspanins associate directly with a small number of primary partners and only indirectly with the wider web, generating a hierarchical rather than a homogeneous network^35,36^, so two tetraspanins with opposing functions are expected to anchor separate neighborhoods rather than co-vary. This supports the TEMD concept - that these tetraspanins act as organizers of compositionally distinct membrane neighborhoods - as a useful framework for circulating myeloid cells, not only endothelium.

Interestingly, CD63 showed the largest change in the family. It is a membrane protein of lysosomes and neutrophil azurophil granules^37^, and granule-protein transcripts are a hallmark of the immature neutrophils released by emergency granulopoiesis^38^, so a whole-blood CD63 increase of this size is consistent with the left-shifted neutrophil signature that single-cell studies link to sepsis immunosuppression and to the poor-outcome SRS1 endotype^39^, and whole-blood CD63 correlated with those transcripts here. The single-cell data add a second, monocyte source: CD63 was induced within CD14 and CD16 monocytes in all three datasets. CD63 is also a surface marker of platelet activation^40^ and the cofactor that retains P-selectin on activated endothelium^41^, but platelet transcripts did not account for the blood signal. CD63 is also the most widely used marker of exosomes and other small extracellular vesicles^42^, so increased cellular CD63 may accompany altered vesicle production or release during inflammation, although transcript abundance does not by itself indicate vesicle secretion. Neutrophils are absent from the PBMC single-cell data, so the CD82 program is characterized only among mononuclear cells. In mouse lung, Cd82 was expressed in roughly 40% of neutrophils and unchanged by LPS. Conversely, the down-regulated block maps onto lymphoid and plasmacytoid dendritic cells depleted in sepsis: TSPAN13 and TSPAN3 are enriched in plasmacytoid dendritic cells and naive B cells^43^, and loss of circulating plasmacytoid dendritic cells predicts death in septic shock^44–46^. CD81, the most strongly down-regulated tetraspanin and the hub of the lymphocyte module, attenuated after composition adjustment and likely reports the same lymphocyte loss. Whether these tetraspanins report only the loss of those cells, or also a change within them, cannot be resolved without neutrophil- and dendritic-cell-inclusive single-cell data.

We find the cell-lineage axis of tetraspanin genes aligns with recent consensus frameworks for sepsis transcriptomics, whereby transcriptional subtypes map to immature neutrophils, platelets and lymphocytes^47^, and the Hi-DEF framework scores immune dysregulation along a myeloid axis with detrimental and protective components versus a lymphoid axis that is protective only^48^. The tetraspanins followed the same division: Myeloid-biased CD63 and CD151 and monocyte-induced CD82 increased and lymphoid-biased TSPAN32 and TSPAN18 decreased in sepsis, and among mortality associations, lymphoid-biased CD37 was associated with survival whereas CD82 was associated with death in the GSE185263 intensive-care arm. This may explain why single-gene mortality associations were frequent but not specific, as most tetraspanins detectable in blood are lineage-dependent. Hi-DEF scores are prognostic after adjustment for the neutrophil-to-lymphocyte ratio^48^, as the CD37 association persisted after composition adjustment here. Whether any tetraspanin adds prognostic information beyond these axes remains to be tested.

These findings may also be relevant to studies of circulating extracellular vesicles (EVs), which are commonly captured and characterized using CD9, CD63 and CD81^42^. In plasma, these tetraspanins mark distinct vesicle subpopulations, with a substantial platelet contribution^49^. If the lineage shift observed here extends to released vesicles, tetraspanin-based capture from septic plasma may enrich for CD63-positive, myeloid-derived vesicles and under-sample CD81-positive vesicles relative to controls, independent of any change in vesicle release per cell. Plasma exosomal CD63 rises from controls to sepsis to septic shock and is associated with mortality^50^, and the monocyte CD63 induction shown here is a candidate cellular source. Endothelial cells are another: CD63 is a component of Weibel-Palade bodies^51^, whose stimulated exocytosis releases CD63-positive intraluminal vesicles^52^, a vascular contribution that blood transcriptomes cannot capture.

Mendelian randomization returned a bounded null: genetically proxied baseline blood tetraspanin expression did not measurably alter sepsis susceptibility. This is consistent with the tetraspanin reorganization we observe being a response to the septic state rather than a germline determinant of who develops sepsis, although a bounded null cannot by itself establish that. The test also addresses susceptibility rather than mortality.

Whereas the only prior tetraspanin-focused sepsis transcriptomic analysis identified erythrocyte-related markers from bulk clustering without survival or cell-type analysis^10^, we add survival analysis, cell-type resolution, composition adjustment, network context and explicit null benchmarking. Because CD82-associated leakage depends on accessible cholesterol and ORP-mediated transport, statins or ORP inhibitors^7^ are testable modulators of the myeloid CD82 program identified here. In the single-cell data, CD151 was detectable in 58% of platelets and CD82 in 19% (46-72% and 9-22% across the three datasets), so mainly the CD151 arm is measurable in platelets. TSPAN9, a platelet-enriched tetraspanin that regulates the collagen receptor GPVI^53,54^, was also modestly up-regulated in septic blood (g=+0.48). Platelets also reorganize their transcriptome and actively translated mRNA pool in sepsis, up-regulating the integrin subunit ITGA2B (αIIb) in a manner that correlates with mortality^55^. Because αIIbβ3 is a CD151-regulated complex, platelets are a plausible next cell type for cell-resolved tetraspanin work.

### Limitations

Cohorts differ in platform, population and sepsis definition, and random-effects modeling and adult-only versus pediatric-only sensitivity analyses mitigate but do not remove this heterogeneity. Time-to-event survival modeling, clinical-covariate adjustment and network analyses rest on a single cohort (GSE65682), whose deposit carries no severity scores, so clinical adjustment was limited to the demographic and diagnostic variables available, and residual confounding by severity cannot be excluded. The single-cell datasets differ in size (2, 7 and 9 healthy donors) and outcome was deposited with only the smallest; the largest shares donors with a bulk cohort in the meta-analysis^25^. The single-cell data also profile PBMCs, so neutrophils - the most expanded cell type in sepsis, expressing CD82 and CD63 - are absent. Deconvolution estimates are reference-based approximations. Two of the eight cohorts are RNA-seq while six are microarray, so platform and biology cannot be fully separated for genes with high between-cohort heterogeneity such as TSPAN2, whose direction differs between cohorts (I²=96%). Finally, the negative ratio result is specific to whole blood and does not exclude a prognostic role for cell-type-specific CD82 or CD151 measurements.

## Conclusions

In human sepsis, tetraspanin expression changes in blood are organized by cell type. The tetraspanins that change appreciably are those abundant in leukocytes, and their direction largely follows myeloid versus lymphoid bias and the compositional shift of sepsis. Two cell-type-specific programs emerge: monocyte CD82 induction marks critical illness, most strongly sepsis, but not consistently outcome, and pan-leukocyte CD37 loss is associated with mortality across ICU cohorts. CD63, the most strongly up-regulated tetraspanin, correlated with immature-neutrophil transcripts and was induced within monocytes. As the canonical exosome marker, CD63 is a candidate link between these leukocyte changes and circulating vesicles. The CD82/CD151 antagonism that motivated the study was established in endothelium, but in blood the two genes change in different cell types and are not transcriptionally coupled, so whole-blood studies cannot assess it. Cell-resolved analyses will therefore be necessary to better understand their mechanistic roles.

## Declarations

### Data availability

All datasets are publicly available from GEO under the accessions listed in Methods. Analysis code is available at https://gitlab.com/jdwren/sepsis_tetraspanin2.

### Author contributions

JDW conceived and implemented the computational analyses in the paper to address the questions and hypotheses to be tested, and drafted the manuscript; YD generated and analyzed the LPS-challenged mouse-lung single-cell RNA-sequencing data; XAZ provided the conceptual foundation and rationale for the study. All authors reviewed, edited and approved the final manuscript.

## Acknowledgements

The authors thank Hunter Porter for helping with manuscript & code revisions.

## Ethics approval

This study analyzed only publicly available, de-identified datasets and did not require new ethics approval.

