## Supplementary tables & figures for "Tetraspanin changes in septic blood are organized by cell type, with monocyte CD82 induction and leukocyte CD37 loss associated with mortality"

### Supplementary Figures

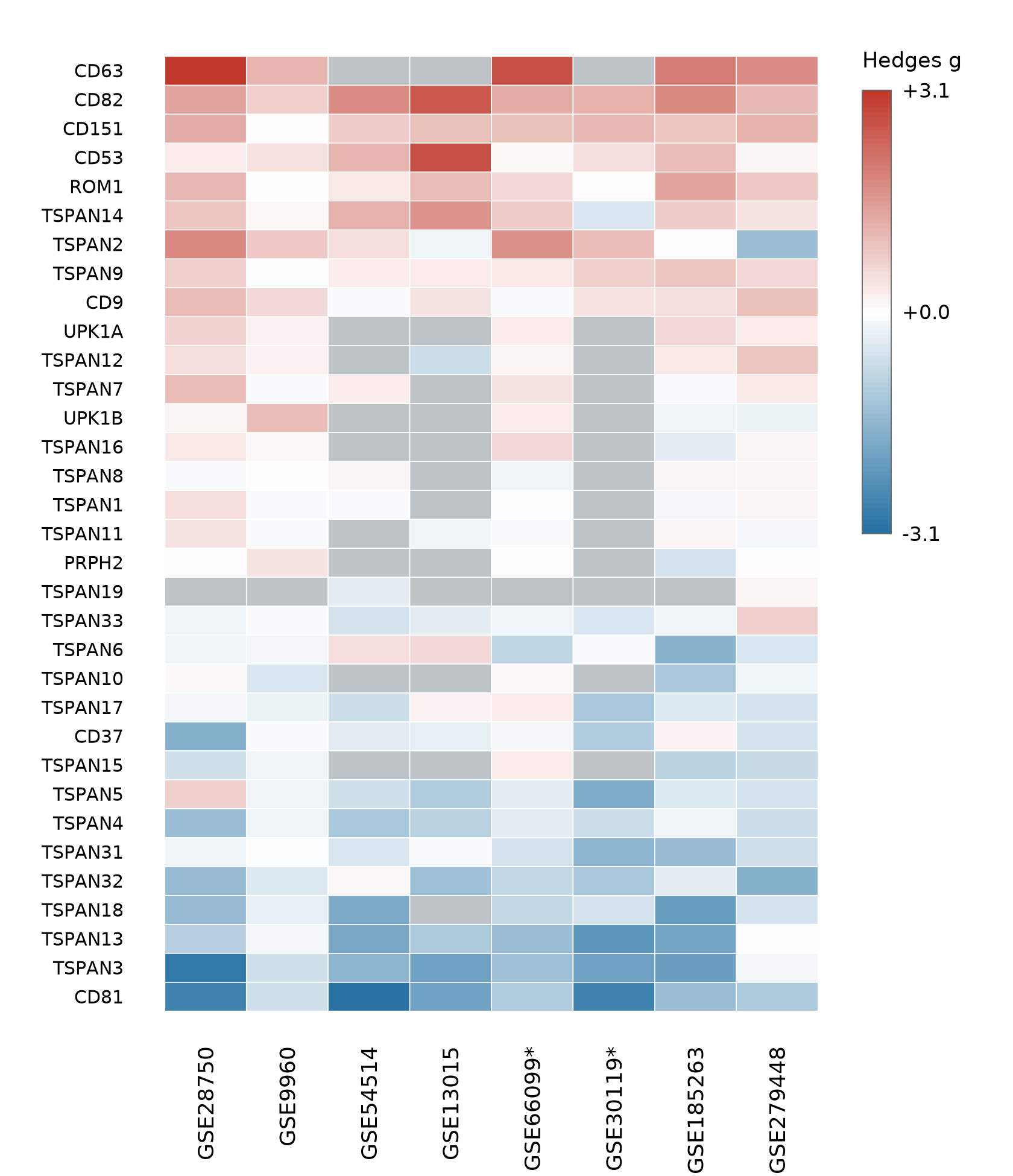

**Supplementary Figure S1.** Heatmap of per-cohort Hedges g (sepsis vs control) for the analyzed tetraspanin genes across the eight ICU-sampled cohorts (asterisk = pediatric cohort; grey = gene not represented on the array), ordered by pooled effect size. The figure shows the per-cohort estimates underlying Figure 1.

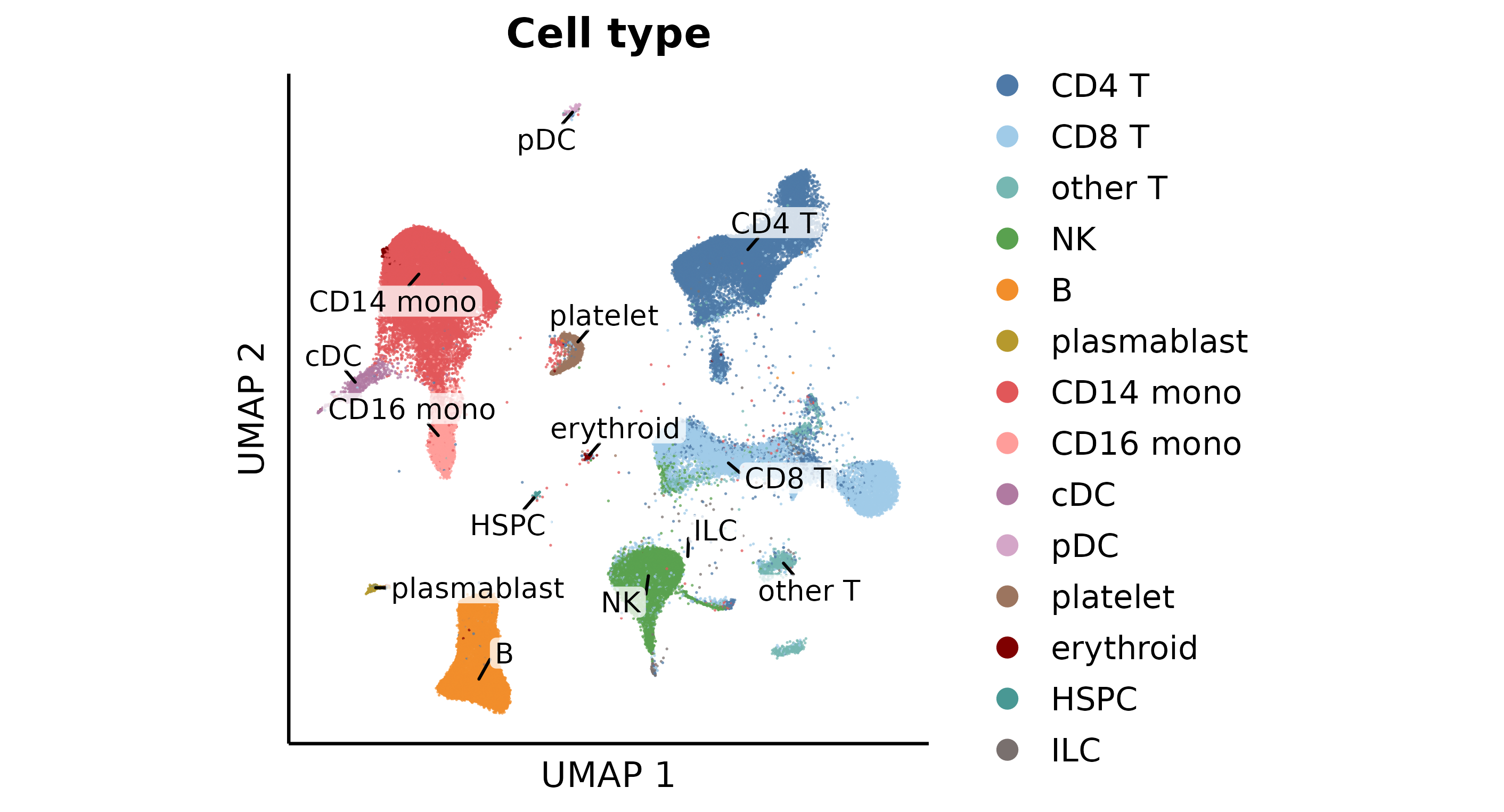

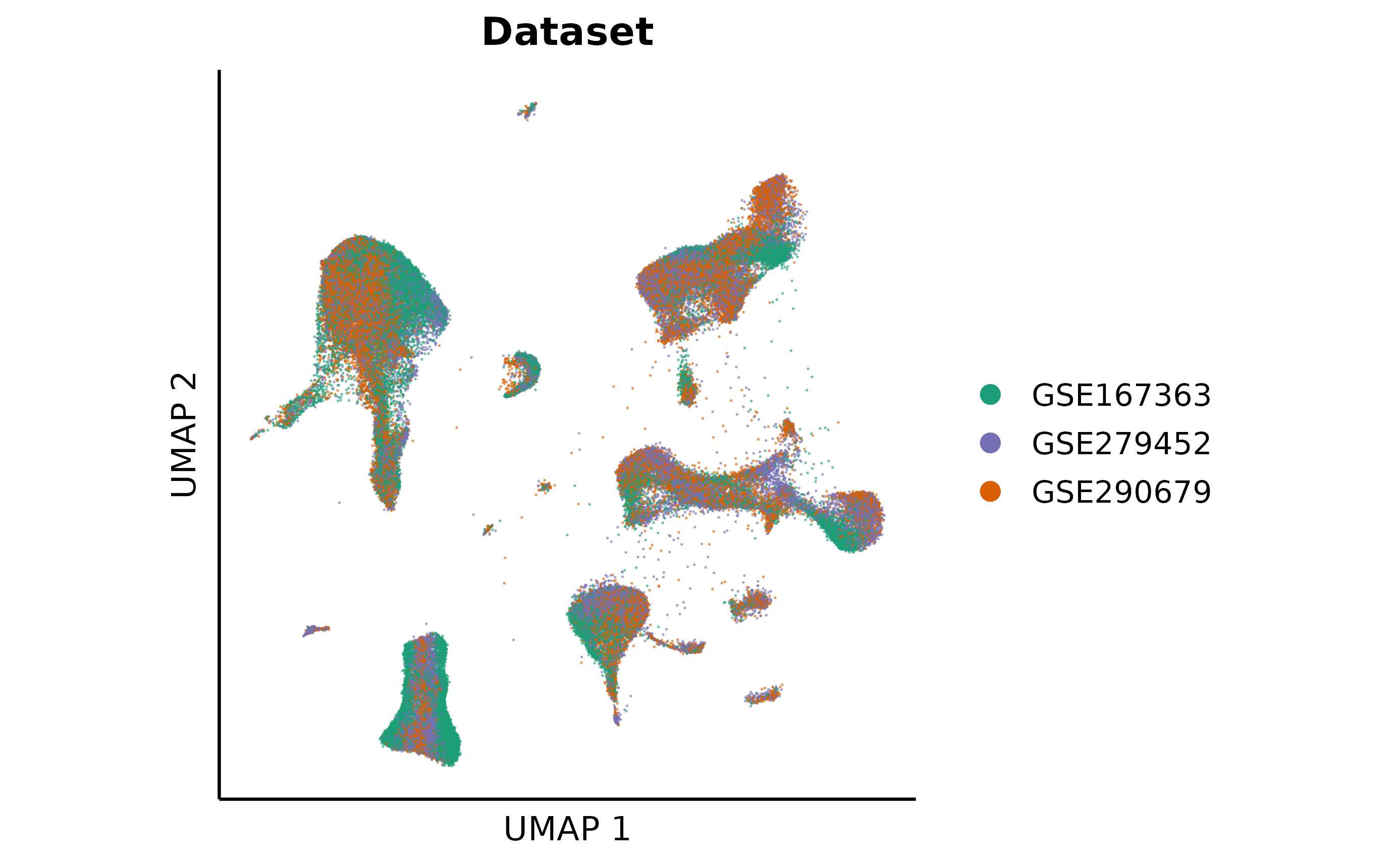

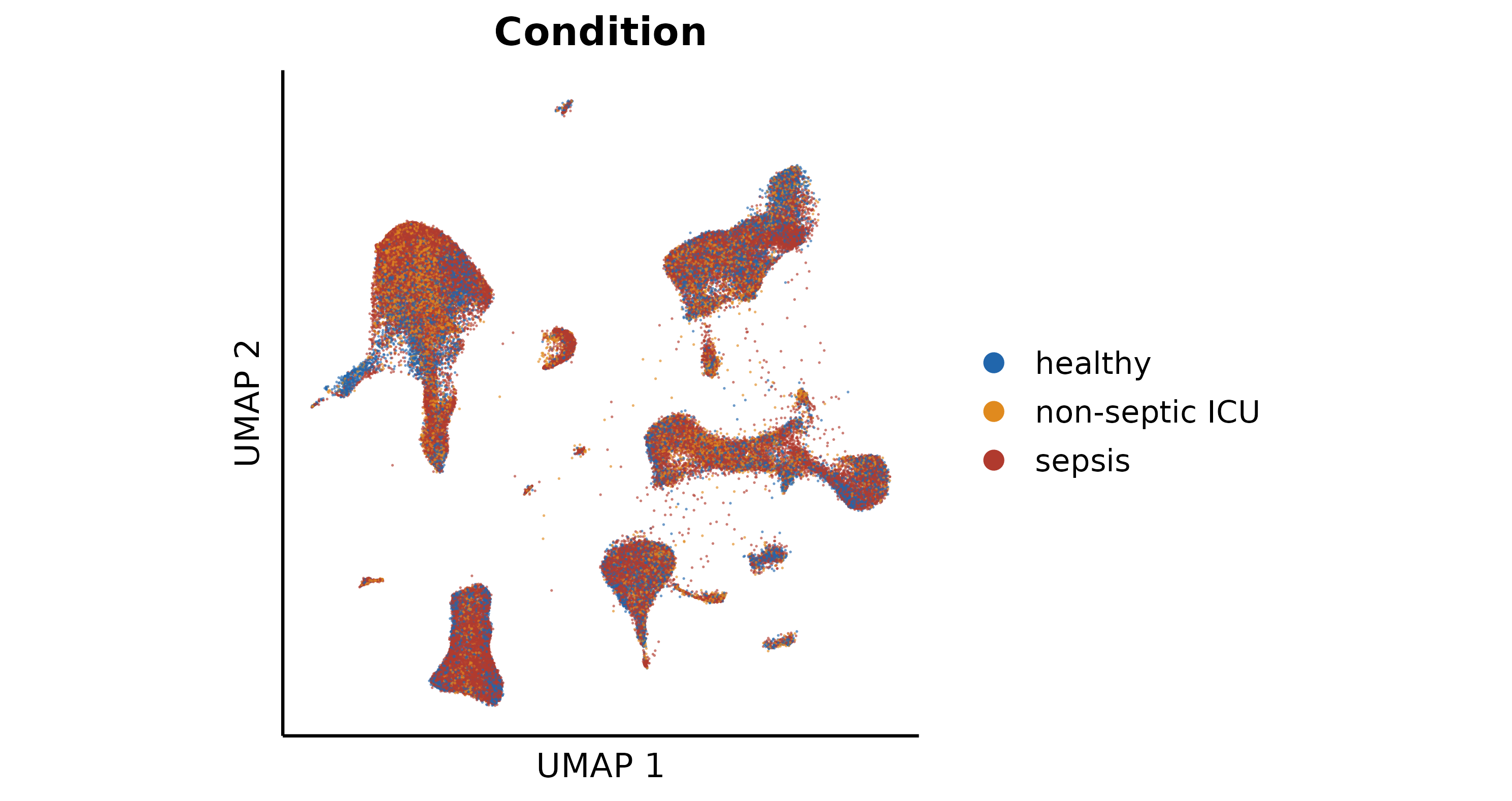

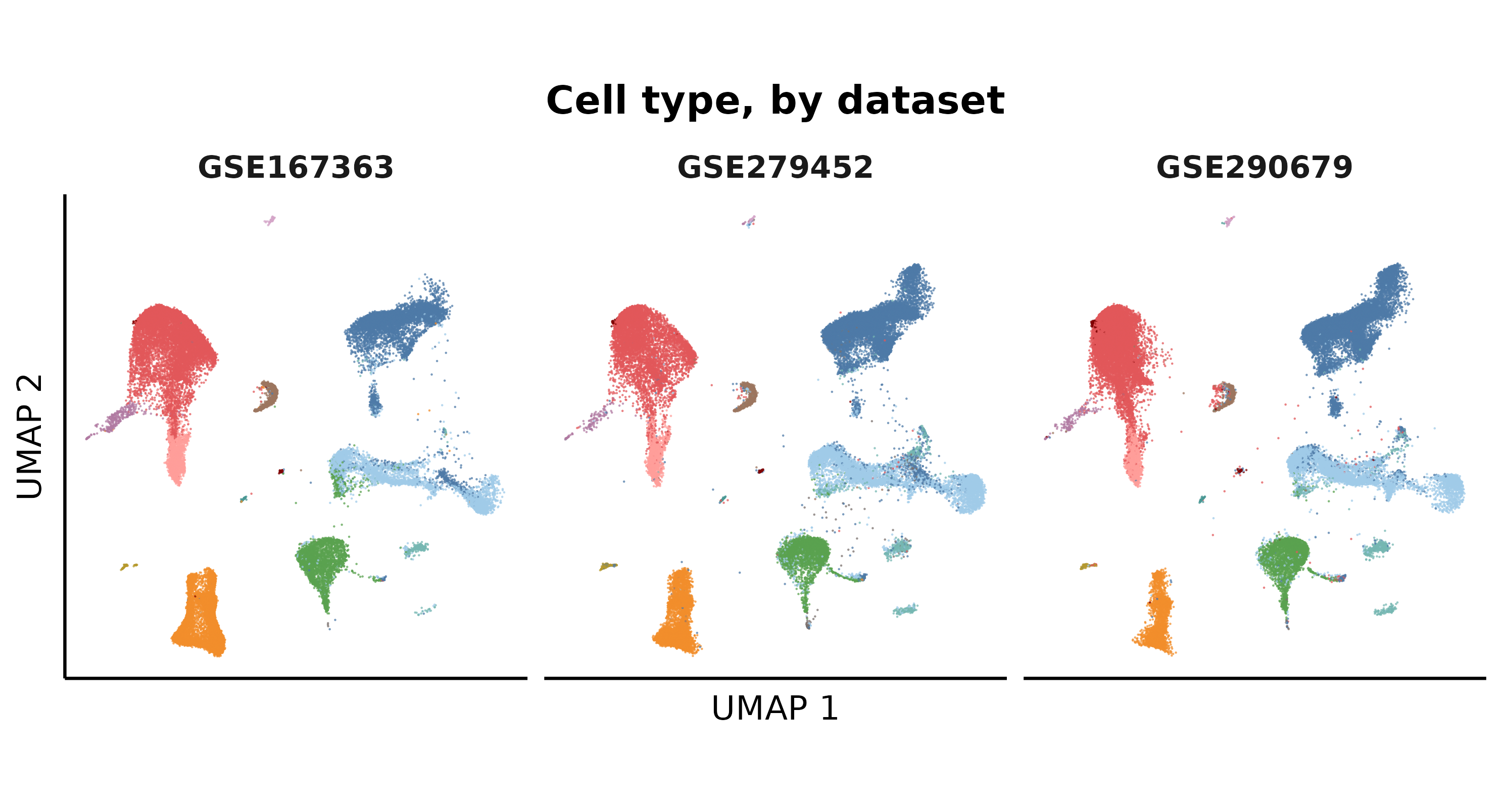

**Supplementary Figure S2.** UMAP of 1,604,899 peripheral-blood mononuclear cells from the three single-cell datasets (189 samples), each cell placed on the Azimuth PBMC reference map (50,000 randomly drawn cells per dataset are displayed), shown as four panels colored by (a) annotated cell type, (b) dataset, (c) condition (healthy, non-septic ICU, sepsis) and (d) annotated cell type with each dataset drawn separately. These annotations underlie the cell-type-resolved differential expression in Figure 4.

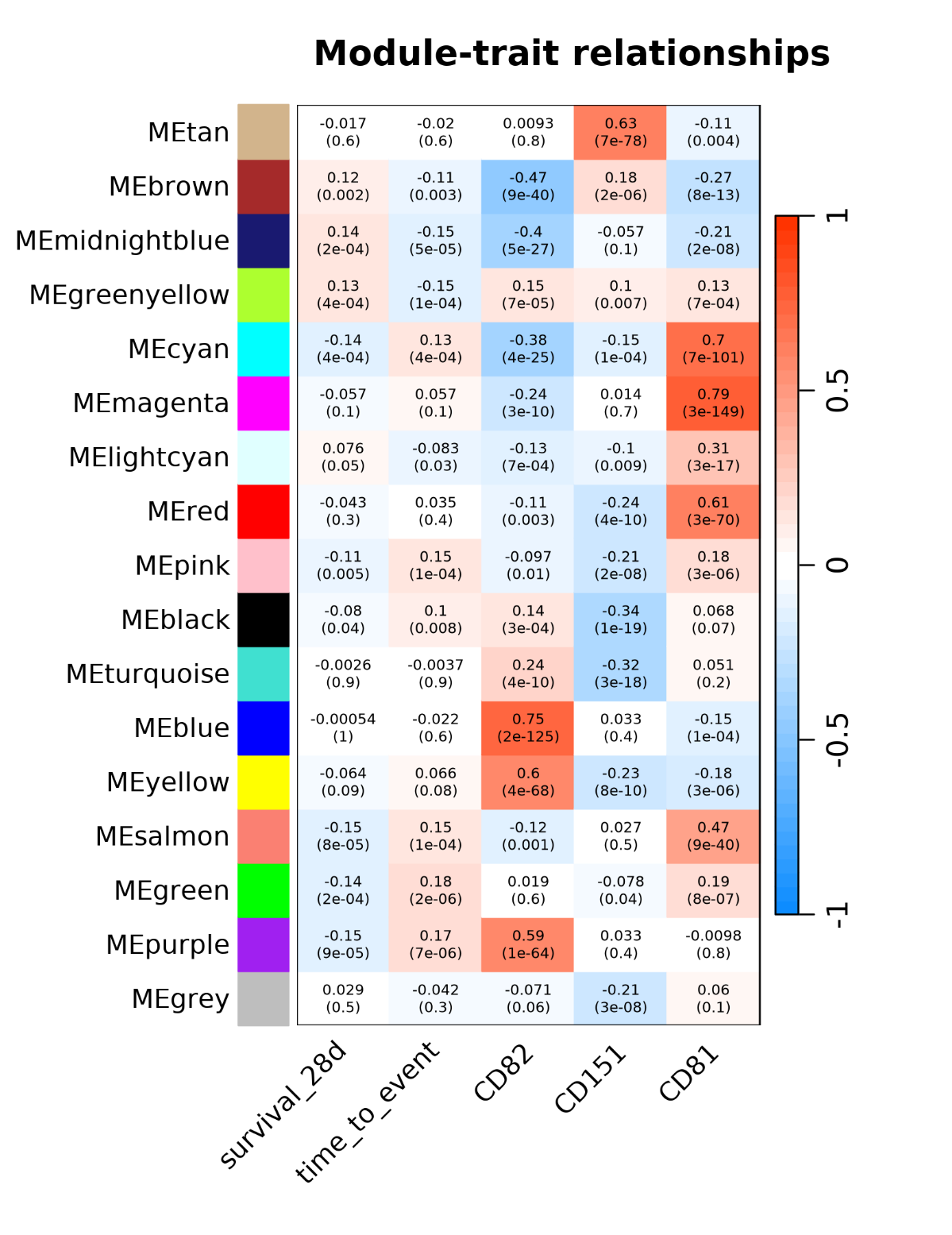

**Supplementary Figure S3.** WGCNA module–trait relationships in 685 intensive-care patients (GSE65682; signed network, soft-threshold power 14). Each cell shows the Pearson correlation (with p-value) between a module eigengene and 28-day survival, time-to-event, or CD82/CD151/CD81 expression. CD82 correlates most strongly with the blue (r = 0.75) and yellow (r = 0.60) modules, CD151 with the tan module (r = 0.63), and CD81 with the magenta module (r = 0.79); module colors are arbitrary labels assigned by WGCNA in order of module size, supporting the module assignments reported in the text.

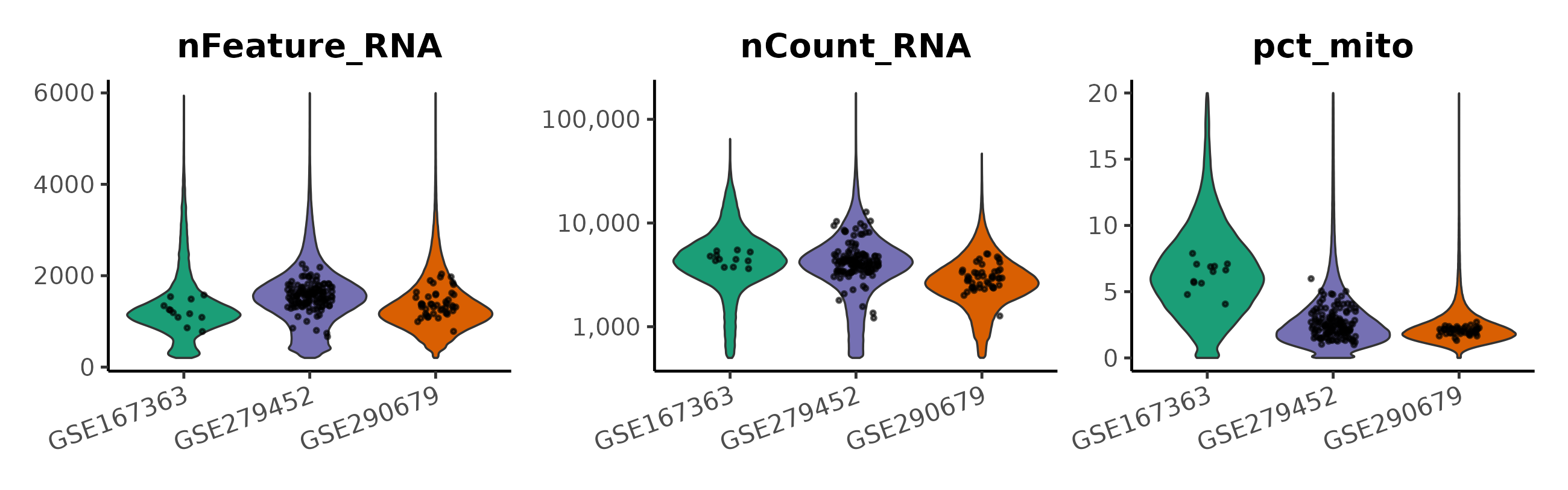

**Supplementary Figure S4.** Per-sample single-cell quality-control distributions for the three single-cell datasets (189 samples; points are per-sample medians, unique transcripts on a log scale): detected genes (nFeature_RNA), unique transcripts (nCount_RNA) and mitochondrial content (pct_mito) after filtering (200–6,000 detected features, <20% mitochondrial reads) and doublet removal.

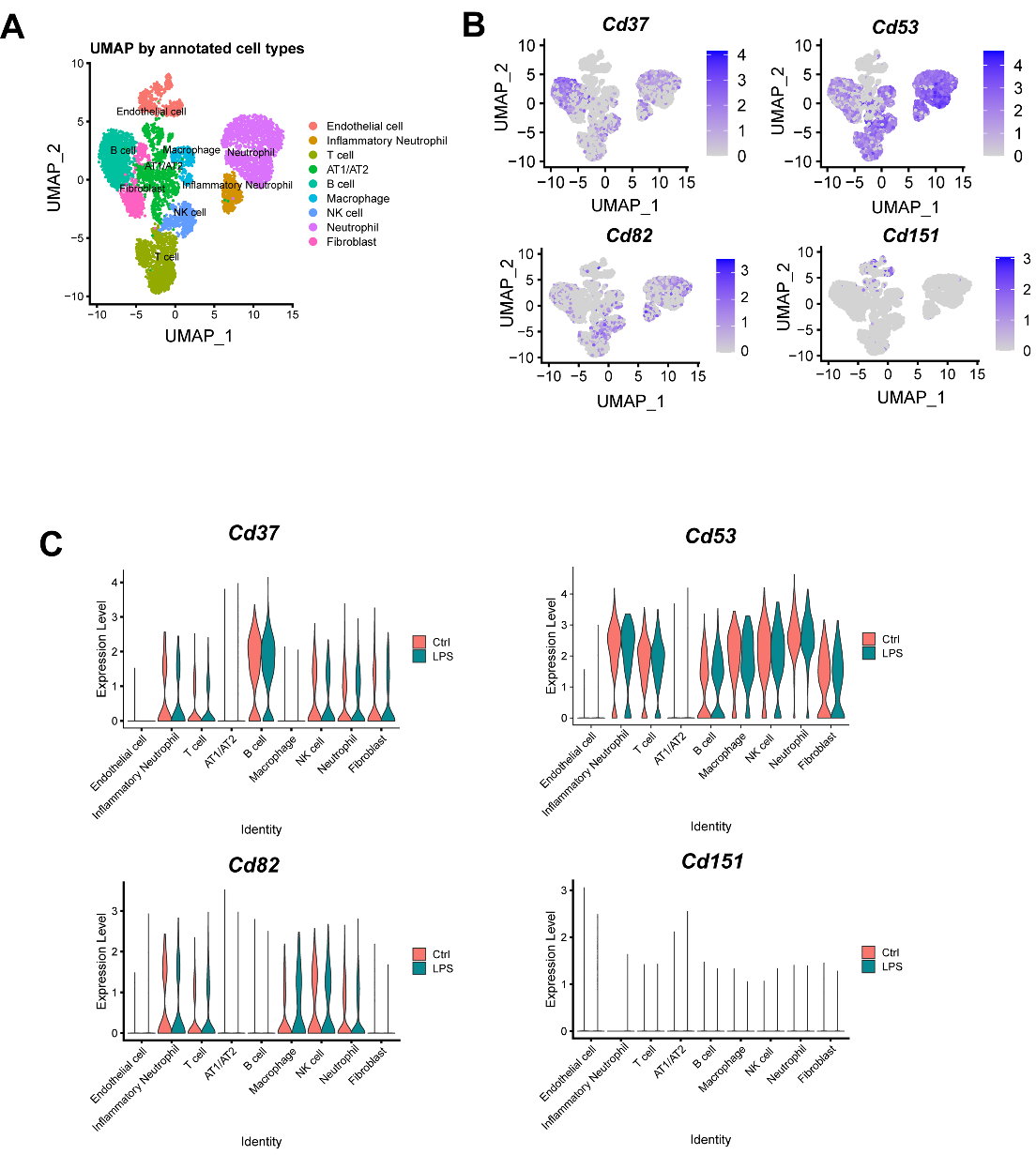

**Supplementary Figure S5.** Single-cell RNA-sequencing of mouse lung following LPS challenge. (A) UMAP of annotated pulmonary cell populations. (B) Feature plots and (C) split violin plots (Ctrl versus LPS) of Cd37, Cd53, Cd82 and Cd151 by population. Cd151 is detected in roughly 30 to 50 percent of endothelial cells and is near-absent (<5%) from the other eight populations.

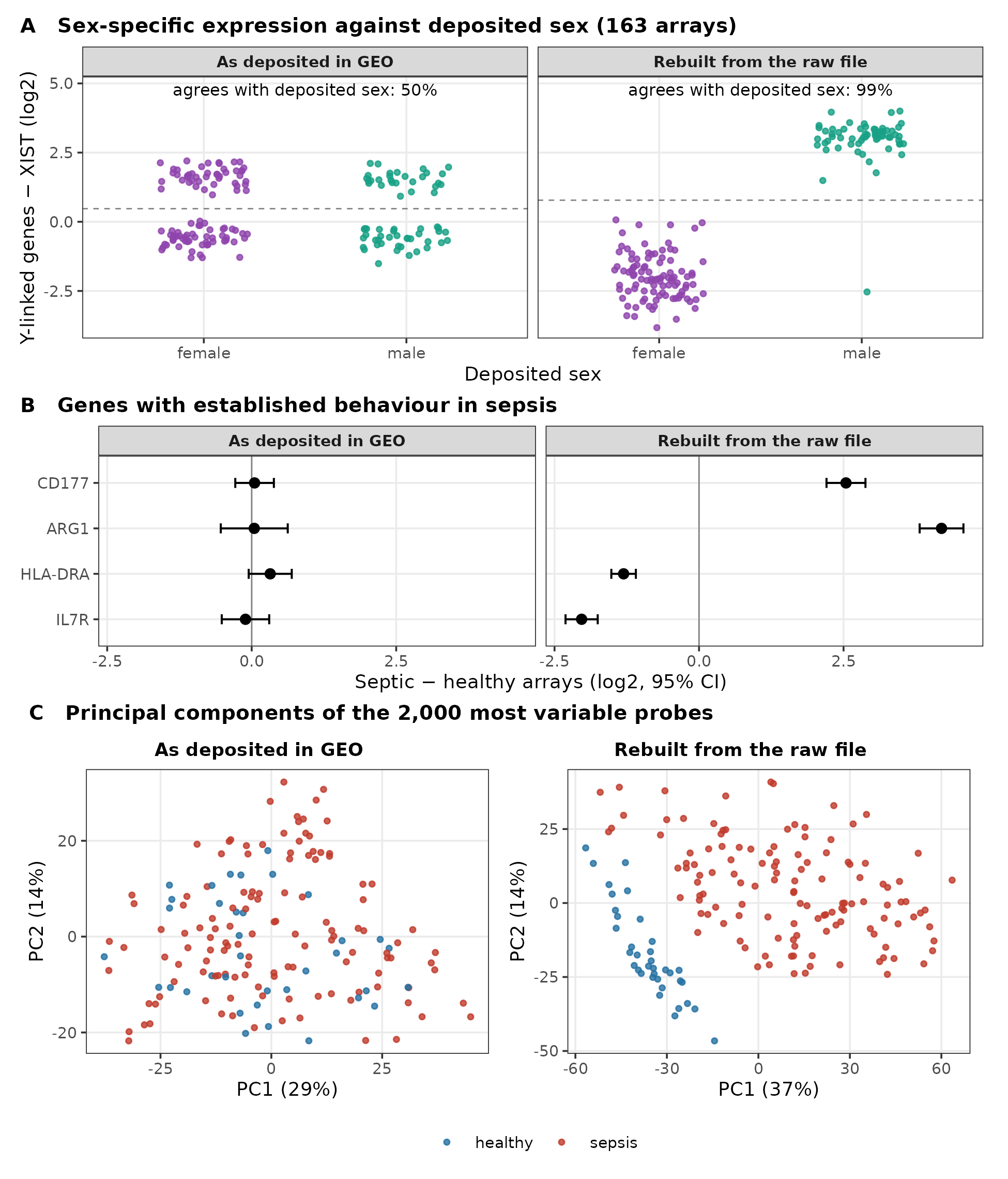

**Supplementary Figure S6.** GSE54514 as deposited in GEO and as rebuilt from its raw file. (A) Sex-specific expression (mean of RPS4Y1, DDX3Y, KDM5D and EIF1AY minus XIST) by deposited sex for the 163 arrays - the dashed line marks the largest gap in the score. Deposited sex agrees with expression in 50% of arrays as deposited and in 99% after rebuilding, and 35 of the 54 participants change expressed sex across their serial arrays as deposited, vs 1 after. (B) Difference between septic and healthy arrays (log2, 95% CI) for four genes with established behavior in sepsis (CD177 and ARG1 induced, HLA-DRA and IL7R reduced). (C) First two principal components of the 2,000 most variable probes. Healthy and septic arrays separate along the first component only after rebuilding (area under the curve 0.95, against 0.55 as deposited).

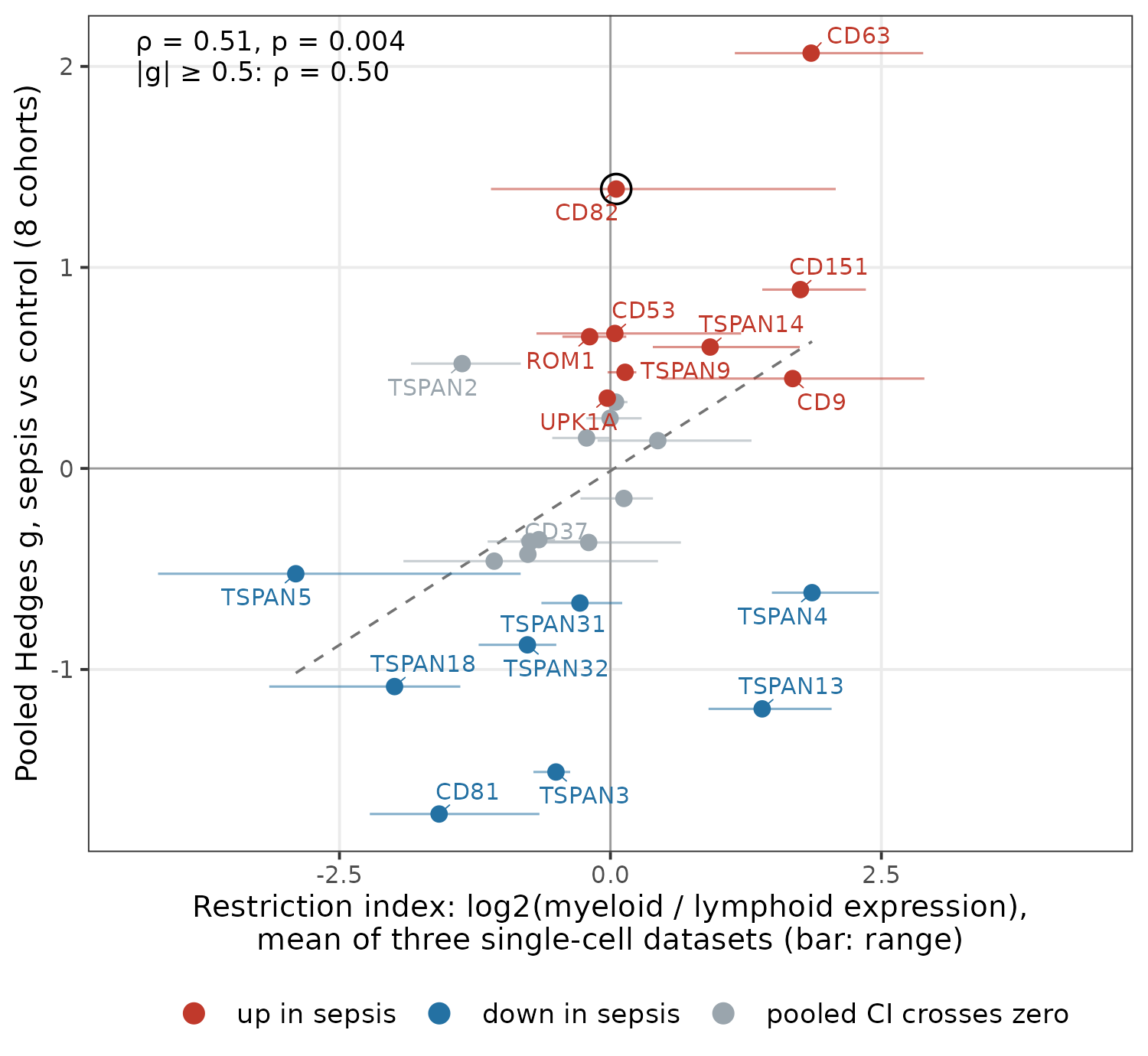

**Supplementary Figure S7.** Myeloid/lymphoid restriction index against the pooled sepsis effect. Each point is one of the 28 tetraspanins in the single-cell analysis. x-axis, log2 ratio of mean expression in monocytes and dendritic cells to that in T, B and NK cells, averaged over the three single-cell datasets, with a horizontal bar spanning the three individual values; y-axis, pooled Hedges g from the eight-cohort meta-analysis (Figure 1). Red, increased in sepsis; blue, decreased; grey, pooled 95% CI crosses zero. Spearman ρ=0.51 (one-sided permutation p=0.004) across all genes and ρ=0.50 among the 15 with |g|≥0.5; the values for each dataset are given in the text. The dashed line is a least-squares fit for orientation. CD82 (circled) has a positive index only in GSE167363.

| **Gene** | **cis-eQTL instrument (eQTLGen)** | **Analyzed in MR** | **SMR p (BH q)** | **Coloc PP.H4** | **Notes** |
| --- | --- | --- | --- | --- | --- |
| **TSPAN13** | Yes | Yes | 0.009 (0.20) | 0.14 | - |
| **CD63** | Yes | Yes | 0.016 (0.20) | 0.085 | - |
| **CD53** | Yes | Yes | 0.050 (0.39) | 0.032 | - |
| **TSPAN2** | Yes | Yes | 0.065 (0.39) | 0.027 | - |
| **CD82** | Yes | Yes | 0.153 (0.64) | 0.014 | - |
| **TSPAN33** | Yes | Yes | 0.159 (0.64) | 0.012 | - |
| **CD9** | Yes | Yes | 0.319 (0.98) | 0.008 | - |
| **PRPH2** | Yes | Yes | 0.416 (0.98) | 0.007 | - |
| **TSPAN10** | Yes | Yes | 0.490 (0.98) | 0.007 | - |
| **TSPAN32** | Yes | Yes | 0.497 (0.98) | 0.006 | - |
| **TSPAN14** | Yes | Yes | 0.620 (0.98) | 0.005 | - |
| **TSPAN18** | Yes | Yes | 0.668 (0.98) | 0.006 | - |
| **TSPAN5** | Yes | Yes | 0.702 (0.98) | 0.010 | - |
| **CD81** | Yes | Yes | 0.708 (0.98) | 0.006 | - |
| **TSPAN3** | Yes | Yes | 0.712 (0.98) | 0.005 | - |
| **TSPAN31** | Yes | Yes | 0.721 (0.98) | 0.009 | - |
| **CD37** | Yes | Yes | 0.774 (0.98) | 0.007 | - |
| **TSPAN16** | Yes | Yes | 0.807 (0.98) | 0.006 | - |
| **TSPAN17** | Yes | Yes | 0.858 (0.98) | 0.005 | - |
| **TSPAN15** | Yes | Yes | 0.895 (0.98) | 0.006 | - |
| **TSPAN9** | Yes | Yes | 0.908 (0.98) | 0.010 | - |
| **CD151** | Yes | Yes | 0.953 (0.98) | 0.005 | - |
| **ROM1** | Yes | Yes | 0.974 (0.98) | 0.005 | - |
| **TSPAN4** | Yes | Yes | 0.979 (0.98) | 0.005 | - |
| **UPK1A** | Absent | No | - | - | No cis probe in eQTLGen; GTEx whole-blood sensitivity arm null (PP.H4=0.005) |
| **UPK1B** | Absent | No | - | - | No cis probe in eQTLGen |
| **TSPAN1** | Absent | No | - | - | No cis probe in eQTLGen |
| **TSPAN6** | Absent | No | - | - | No cis probe in eQTLGen |
| **TSPAN7** | Absent | No | - | - | No cis probe in eQTLGen |
| **TSPAN8** | Absent | No | - | - | No cis probe in eQTLGen |
| **TSPAN11** | Absent | No | - | - | No cis probe in eQTLGen |
| **TSPAN12** | Absent | No | - | - | No cis probe in eQTLGen |
| **TSPAN19** | Absent | No | - | - | No cis probe in eQTLGen; also not represented in the expression meta-analysis |

**Supplementary Table S1.** All 33 canonical human tetraspanins and their status in the Mendelian-randomization analysis (eQTLGen whole-blood cis-eQTL exposure to the FinnGen R12 sepsis outcome). Twenty-four genes carried a genome-wide-significant whole-blood cis-eQTL instrument (instrument p≤5×10⁻⁸) and were tested by SMR/HEIDI and colocalization; nine lacked a cis probe in eQTLGen and were not instrumentable. No gene reached Benjamini-Hochberg FDR<0.05 or colocalized (PP.H4≥0.75). Genes are ordered by ascending SMR p among those tested, then by absence. BH q, Benjamini-Hochberg false-discovery rate; PP.H4, posterior probability of a shared causal variant.

| **Series** | **Samples / subjects** | **Draws per subject** | **Platform** | **Controls (samples / subjects)** | **Patient overlap with** | **Used in this study** |
| --- | --- | --- | --- | --- | --- | --- |
| **GSE28750** | 41 / 41 | 1 | GPL570 | 20 / 20 | none | meta, deconv. (20 ctrl, 10 sepsis) |
| **GSE9960** | 70 / 70 | 1 | GPL570 | 16 / 16 | none | meta (16 ctrl, 54 sepsis) |
| **GSE54514** | 163 / 54 | ≤5 (daily) | GPL6947 | 36 / 18 | none | meta, deconv., mortality (D1 only: 18 ctrl, 35 sepsis); rebuilt from raw file |
| **GSE13015** | 106 / 106 | 1 | GPL6106, GPL6947 | 29 / 29 | none | meta (5 ctrl, 29 sepsis; GPL6947 arm) |
| **GSE66099** | 276 / 276 | 1 (ICU D1) | GPL570 | 47 / 47 | its 6 constituent series (below) | meta, deconv. (47 ctrl, 199 sepsis) |
| **GSE30119** | 143 / 143 | 1 | GPL6947 | 44 / 44 | none | meta, deconv. (44 ctrl, 9 sepsis-flagged) |
| **GSE185263** | 392 / 392 | 1 | GPL16791 | 44 / 44 | none | meta, deconv. (44 ctrl, 82 ICU sepsis); mortality (ICU arm; ED arm separate) |
| **GSE279448** | 207 / 207 | 1 | GPL24676 | 75 / 75 | GSE279452 (84 donors) | meta (47 healthy ctrl, 132 sepsis; 28 ICU ctrl excluded) |
| **GSE65682** | 802 / 802 | 1 | GPL13667 | 84 / 84 | none | survival, WGCNA (ctrl not used) |
| **GSE95233** | 124 / 71 | ≤4 (D0–D3) | GPL570 | 22 / 20 | none | external mortality validation (D1) |
| **GSE167363** | 12 / 7 | ≤2 (T0, T6) | GPL24676 | 2 / 2 | none | single-cell (T0) |
| **GSE279452** | 133 / 133 | 1 | GPL24676 | 23 / 23 | GSE279448 (84 donors) | single-cell, adults (7 healthy ctrl, 92 sepsis, 8 ICU ctrl); 1 sex-discordant sample excluded |
| **GSE290679** | 47 / 47 | 1 | GPL24676 | 28 / 28 | none | single-cell (9 healthy ctrl, 19 sepsis, 17 ICU ctrl); 2 sex-discordant samples excluded |
| **GSE4607** | 123 / 83 | ≤2 (D1, D3) | GPL570 | 15 / 15 | GSE66099 | excluded |
| **GSE8121** | 75 / 46 | ≤2 (D1, D3) | GPL570 | 15 / 15 | GSE66099 | excluded |
| **GSE9692** | 45 / 45 | 1 | GPL570 | 15 / 15 | GSE66099 | excluded |
| **GSE13904** | 227 / 152 | ≤2 (D1, D3) | GPL570 | 18 / 18 | GSE66099 | excluded |
| **GSE26440** | 130 / 130 | 1 | GPL570 | 32 / 32 | GSE66099 | excluded |
| **GSE26378** | 103 / 103 | 1 | GPL570 | 21 / 21 | GSE66099 | excluded |

**Supplementary Table S2.** Sample-independence audit of every GEO series used in this study. For each series: total samples, unique subjects, timepoints per subject, platforms, control samples versus control individuals, cross-series patient overlap, and how the series was used here. GSE66099 is a third-party reanalysis compiling the Day-1 unique patients of GSE4607, GSE8121, GSE9692, GSE13904, GSE26378 and GSE26440^15^; its 276 samples are provenance-linked to their originating samples, and its six constituent series are therefore excluded from the meta-analysis and deconvolution reported here. Abbreviations: ctrl, control; D, day; T, time point; ICU, intensive care unit; ED, emergency department; meta, meta-analysis; deconv., deconvolution; n/a, not applicable (series excluded).
